# Tri-omic mapping revealed concerted dynamics of 3D epigenome and transcriptome in brain cells

**DOI:** 10.1101/2024.05.03.592322

**Authors:** Haoxi Chai, Xingyu Huang, Guangzhou Xiong, Jiaxiang Huang, Katarzyna Karolina Pels, Lingyun Meng, Jin Han, Dongmei Tang, Guanjing Pan, Liang Deng, Qin Xiao, Xiaotao Wang, Meng Zhang, Krzysztof Banecki, Dariusz Plewczynski, Chia-Lin Wei, Yijun Ruan

## Abstract

Exploring the genomic basis of transcriptional programs has been a longstanding research focus. Here, we report a high-throughput single-cell tri-omic method to capture <u>ch</u>romatin <u>a</u>ccessibility, interaction, and <u>R</u>NA simultaneously (ChAIR). After validating in cultured cells, we applied ChAIR to brain cells across mouse lifespan and delineated the concerted dynamics of 3D-epigenomic architecture and transcription during maturation and aging. Particularly, ultra-long chromatin megacontacts and promoter-associated 3D-epigenomic states are effective in defining cell identity and revealing spatially-resolved anatomic specificity. Importantly, we found that neurons in different brain regions and non-neuronal cells may undergo divergent genomic mechanisms during differentiation and aging. Our results demonstrated ChAIR’s robustness of connecting chromatin folding architecture with cellular property and its potential applications to address complex questions in single-cell resolution and spatial specificity.

## Introduction

It has been presumed that the three-dimensional (3D) genome folding architecture would profoundly determine genome functions including gene transcription regulation(*1–3*). However, much of the existing insights into this architecture have been gleaned largely from bulk-cell data e.g., ChIA-PET(*4*), Hi-C(*5*), and related methods(*6, 7*), revealing essential but averaged views including chromatin compartments(*8*), topologically associated domains (TADs)(*9*), and chromatin loops(*10*). Meanwhile, recent functional assays, by perturbing chromatin folding structures, showed minor attenuation to gene transcription(*11*), and inversely, disturbing transcription activity resulted in minimal impact on the overall genome topology(*12*), raising puzzlement regarding the true causal relationship of chromatin topology and gene transcription. Although technological advancements in single-cell Hi-C(*13–16*) and Dip-C(*17*) for 3D genome mapping have illuminated the stochastic characteristics of chromatin interactions and unveiled detailed chromosome structures through 3D modeling within individual cells(*13–17*), yet fall short of elucidating the mechanistic links between genome structure and its biological implications. Bridging this gap, multi-omic single-cell methods including scCARE-seq(*18*), HiRES(*19*), LiMCA(*20*) and GAGE-seq(*21*) were developed recently to simultaneously capture chromatin interactions and gene expression (Hi-C+RNA-seq) in the same cells, demonstrating the technical possibility to directly establish the functional relevance of 3D genome structure to gene transcription during lineage specification in single cells. Still, the current single-cell methods with limited modality and low-throughput are not fully equipped to unravel the detailed relationship between epigenomic states, 3D genome folding architecture, and transcription regulation concomitantly. Here, we introduce a tri-omic mapping assay, ChAIR, which simultaneously maps chromatin accessibility, interactions, and RNA in individual cells. After validating this method with cell-cycle specific profiles of cultured cell lines, we applied ChAIR to mouse brain cells across 5 age points of lifespan, capturing the concurrent changes in 3D genome folding, epigenome remodeling, and transcription modulation during cell maturation and aging. Furthermore, by integrating ChAIR data with spatial transcriptomic data, we established a strategy to unveil spatially-resolved 3D epigenomic features (3D genome and epigenome) with anatomic specificity. Specifically, the tri-omic ChAIR data enabled us to unravel the genomic features of chromatin interactions and open chromatin loci that were uniquely associated with the promoters of cell-type specific genes in different brain cells. Notably, the dynamic balance of ultra-long chromatin contacts in single cell genomes appeared to be a valuable indicator of transitions in brain cell lineage and aging processes. We also found that neurons appear to employ different genomic mechanisms compared to non-neuronal cells during cell differentiation. Meanwhile, neurons from different brain regions exhibit divergent strategies in adapting to aging. Through these analyses, we demonstrated ChAIR’s ability to capture the direct correlation between genomic architectures, epigenomic states, and gene expression together in single cells, and revealed novel insights into the genomic basis of brain cell maturation and aging.

## Results

### ChAIR captures tri-omic data in single cells

In order to develop a single cell method to simultaneously capture chromatin interactions and epigenomic states, we have previously established a bulk-cell dual-omic method called ChIATAC(*22*). ChIATAC captures open chromatin loci similar to ATAC-seq(*23*) and chromatin interactions between those loci simultaneously. By adopting ChIATAC to a microfluidic system, we developed a protocol to capture single-cell <u>Ch</u>romatin <u>A</u>ccessibility, Interaction and <u>R</u>NA (called ChAIR) simultaneously in same individual cells. This procedure involves crosslinking cells, isolating and permeabilizing nuclei followed by *in situ* procedures that include chromatin DNA restriction digestion, proximity ligation, and Tn5 tagmentation. Then, the intact nuclei were uploaded into the 10x Genomics Multiome system, capturing barcoded DNA and RNA molecules simultaneously in single cells, and the corresponding DNA and RNA information were revealed by sequencing (Fig. 1A). Each ChAIR experiment generates three datasets (tri-omics) including ChAIR-ATAC data for chromatin accessibility, ChAIR-PET (paired-end-tag) for chromatin interaction, and ChAIR-RNA for gene expression. Thus, this tri-omic approach provides a unique strategy to investigate the intricate interplays of epigenome, 3D genome and transcriptome directly in the same individual cells in a high-throughput manner.

**Fig. 1.**
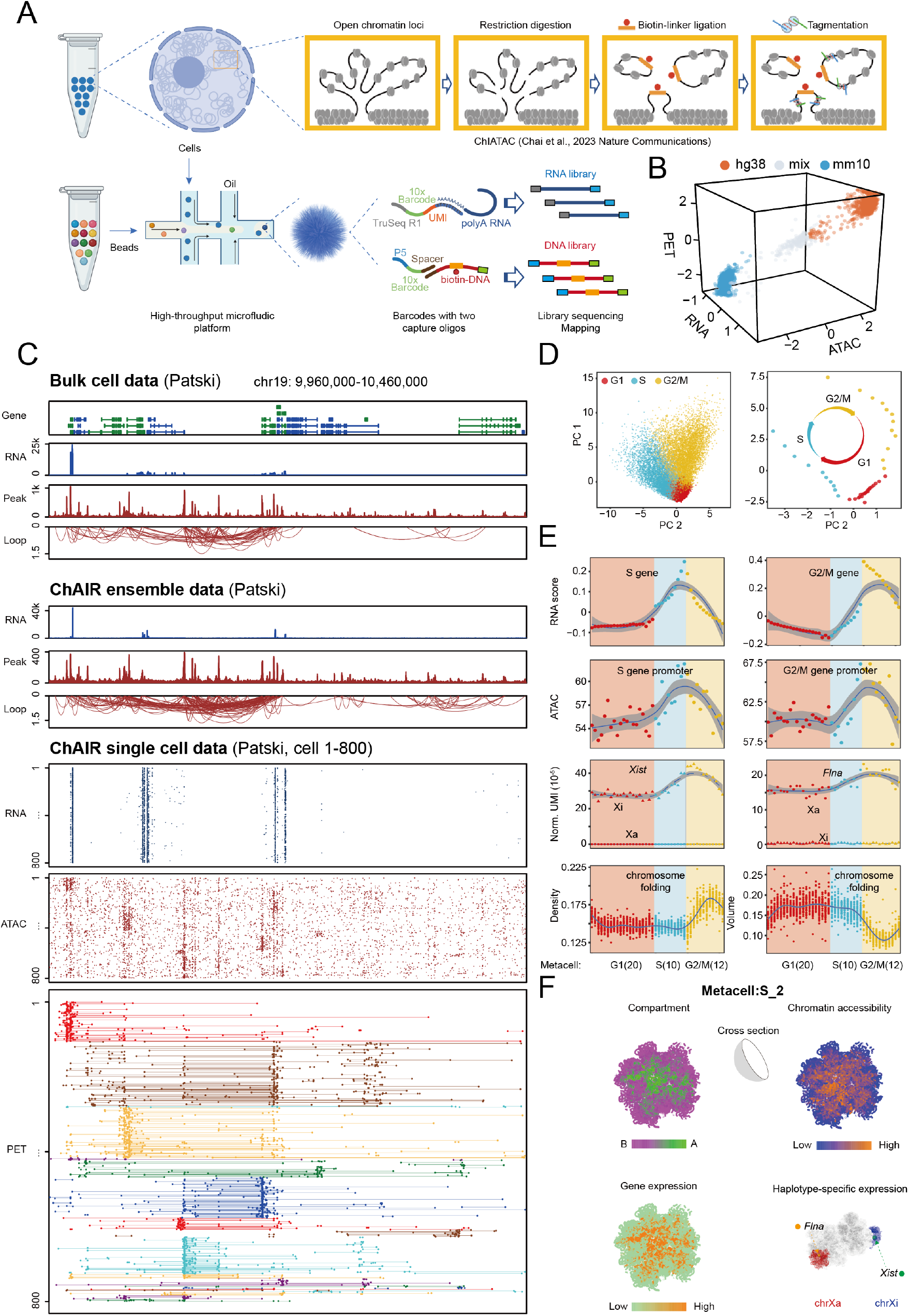
ChAIR captures comprehensive 3D epigenome and transcriptome landscapes in single cells. (**A**) Schematics of ChAIR. (**B**) A 3D scatter plot of ChAIR tri-omic data (RNA, ATAC, PET) in a Barnyard experiment using mixed K562 and Patski cells. (**C**) Browser-based visualization of Patski bulk-cell ChIATAC and RNA-seq data, ChAIR data in ensemble, and in single-cell tracks. (**D**) UMAP of Patski ChAIR-RNA data with cell cycle-specificity (left) and 42 metacells ordered by cell cycle pseudotime (right). (**E**) Characterization of cell cycle-specific and haplotype-resolved Patski ChAIR-RNA data (RNA score) for S-specific genes (S gene) and G2/M-specific genes (G2/M gene), ChAIR-ATAC data at promoters of S genes and G2/M genes, normalized UMI counts of *Xist* and *Flna* expression in chrXi and chrXa, along with estimates of chromatin folding density and nuclear volume using ChAIR-PET data across 42 metacells. (**F**) Example views of 3D models of nuclear architectures with chromatin compartment A and B, open chromatin accessibility, gene expression, and haplotype-resolved nuclear positioning of *Xist* and *Flna* in chrXa and chrXi.

Through a proof-of-concept experiment using an equal mixture of human K562 (a myelogenous leukemia cell line) and mouse Patski (a hybrid mouse fibroblast cell line) cells, we demonstrated the effectiveness of the ChAIR protocol for generating high quality tri-omic datasets with an overall 5.2% doublet rate (Fig. 1B and fig. S1A), confirming the authenticity of our microfluidic system for single-cell analysis. With meticulous optimization, we have successfully established a robust ChAIR protocol and generated single-cell data from 35,515 Patski cells and 40,595 K562 cells (table S1, Fig. 1C and fig. S2 A-B) with high reproducibility (fig. S1B-C). Compared to single-cell data derived from K562 cells(*24–26*), the ChAIR-RNA and -ATAC data showed a similar level of robustness in capturing gene transcripts and mapping accessible chromatin sites (fig. S3A-D). While the total number of chromatin contacts per cell in ChAIR-PET data was lower compared to the mono-modal sci-Hi-C data(*27*), the number of TSS-associated contacts was comparable (fig. S3E). Furthermore, the ratios of TSS-associated chromatin contacts over the non-TSS contacts in ChAIR-PET data were significantly higher than that in sci-Hi-C data (fig. S3F). This result underscores ChAIR’s effectiveness in reducing random background noise and enriching chromatin interactions at open chromatin loci and transcription active regions.

The ensemble ChAIR data analysis showed expected characteristics in comparison with the bulk cell data (Fig. 1C). The majority of the ChAIR-RNA reads were uniquely mapped to exon regions (fig. S4A) similar to the bulk-cell RNA-seq data. Additionally, the ensemble ChAIR-ATAC data exhibited the enrichment at open chromatin loci and transcription start sites (TSSs) comparable to those observed in 10x Multiome ATAC, ChIATAC and ATAC-seq, whereas the Hi-C data showed no signal enrichment (fig. S4B). Chromatin state analysis by ChromHMM(*28*) also showed that the ChAIR-identified open chromatin loci and chromatin loops shared the same epigenomic property as demonstrated in bulk-cell ATAC-seq and ChIATAC data, specifically showing enrichment for promoter-associated characteristics (fig. S4C-D). In addition, the ensemble ChAIR-PET data and the ChIATAC data shared the same profile for the lengths of chromatin loops with various interaction frequencies as shown in both K562 and Patski cells (fig. S4E). Additionally, the 2D contact profiles (fig. S5A-D) and the browser-based loop/peak views also showed high similarity between ensemble ChAIR and ChIATAC data (Fig. 1C and fig. S2B).

To visualize the ChAIR data of individual cells, we developed a browser-based visualization tool (see methods), by which the ChAIR data were presented in three tracks (RNA, ATAC, and PET) in alignment with the reference genome (Fig. 1C and fig. S2B). Particularly in the ChAIR-PET data, depicted as two dots connected by a horizontal line, hierarchical clustering revealed that cells within the same cluster shared similar chromatin folding patterns. Conversely, significant variations were observed among different clusters, highlighting cellular heterogeneity that may be attributed to different cell cycle phases. Overall, the ChAIR data showed high efficiency in unveiling the coordinated patterns for gene expression (RNA), open chromatin accessibility (ATAC), and chromatin interactions (PET) in individual cells.

### The interplay between chromatin architecture and transcription during cell cycle

Cell cycle is an important cellular feature leading from DNA replication to cell division. A complete cell cycle includes G1, S, G2, and Mitotic phases. Phase-specific gene transcription has been well characterized(*29*), and chromatin folding architecture during cell cycle has been an active research topic(*30–32*). However, the exact interplays between phase-specific transcriptional programs and chromatin folding dynamics during cell cycle have not been fully characterized. The ChAIR tri-omic data derived from K562 and Patski provided an opportunity to address this question.

We aimed to investigate how chromatin configuration shifts in concert with gene expression changes throughout cell cycle. Using ChAIR-RNA data and the differential expression profiles of S- and G2/M-specific marker genes(*29*) (fig. S6A), we were able to distinguish individual cells representing G1, S, and G2/M phases of Patski and K562, respectively (Fig.1D, left and fig. S6B, left). Analysis of the corresponding ChAIR-PET data revealed high discernibility in compartment strength and TAD signals in G1 and S phases, with a marked decrease in G2/M phase (fig. S6C-D), consistent with findings from prior studies(*30–32*). It is also noticed that the promoter-enhancer interactions (TSS-E) for S-specific and G2/M-specific marker genes showed heightened contact signals during their respective phases (fig. S6E).

To further dissect cell cycle in more details, we ordered the cells in a cell-cycle pseudotime (see methods) based on their positioning on the principal component analysis (PCA) plot, with the trajectory starting from the cells in early G1 phase through S1 and ending in late G2/M phase. Next, we grouped the cells that were proximal to each other on the PCA map into 42 distinct ‘metacells’ (Fig. 1D, right and fig. S6B, right). The cells in each metacell shared highly similar cellular properties, and the chromatin contacts in each metacell provided a robust data representation for analyzing chromatin folding and gene transcription during cellular progression in cell cycle and minimized stochastic fluctuations in individual cells. As expected, the gene expression (RNA) and promoter strength (ATAC) of S-specific marker genes were relatively low in G1, increased upon entry into S, peaked toward the end of S, and declined continuously in G2/M phases, aligning closely with the progression along cell cycle pseudotime (Fig. 1E, top left). Similarly, G2/M marker genes followed this pattern but peaked in mid-G2/M and decreased towards its end (Fig. 1E, top right), demonstrating coordinated changes in gene expression and chromatin accessibility.

Meanwhile, the ChAIR-PET data enabled us to assess cell-cycle dynamics using chromatin contact profiles in each cell based on a well-established method(*16*). As anticipated, five cell-cycle phases (post-M, G1, early S, late S-G2, and pre-M) were identified in K562 and Patski cells, and the majority were in the interphase and particularly in S phase (fig. S7A-B). The intra-chromosomal 2D contact profiles displayed characteristic profiles for cells in different phases, showing marked chromatin compartment and TAD features in interphase and less pronounced structures in mitosis (fig. S7A-B).

Furthermore, because the Patski cell line is derived from a hybrid mouse strain with high frequency of heterozygous SNPs (1 per 70-96 bp)(*33*), we obtained haplotype-resolved ChAIR data and examined the haplotype-specific chromatin folding and gene expression regarding X chromosome inactivation by *Xist* with single-cell resolution during cell cycle. *Xist* was constantly active in chrXi and inactive in chrXa as expected (fig. S8A). From the onset of S phase, the *Xist* activity started to increase, reaching its peak shortly after entering the G2/M phase and then declining in later G2/M phase (Fig. 1E). Conversely, the chrXa-specific gene *Flna*, active on chrXa but not on chrXi (fig. S8B), exhibited fluctuations similar to *Xist*, albeit to a lesser extent, throughout the cell cycle. (Fig. 1E).

Utilizing the 42 metacells along the cell cycle pseudotime of Patski cells (Fig. 1D, right), we explored the haplotype-resolved contact profiles and reconstructed the 3D models of genomes for each of the 42 metacells (Fig. 1F and fig. S9). Comparing the 2D contact heatmaps of metacells corresponding to the G1 and S phases with those in the G2/M phases, we uncovered marked differences in chromatin conformation (fig. S9A-C), in line with previous findings(*16*). This distinct conformational variation was further supported by marked changes in chromatin folding density and volume (see methods) as estimated by 3D modeling(*14*) (Fig. 1E, bottom and fig. S9D-E).

Together, by analyzing ChAIR data from two distinct perspectives, based on either ChAIR-RNA or ChAIR-PET data, our study underscored the synchronized relationship between gene expression, chromatin accessibility, and 3D chromatin conformation throughout the cell cycle, which demonstrated the power of ChAIR to enable efficient single-cell tri-omic analysis for delineating the genomic framework underlying cell-specific transcription regulation.

### Tri-omic landscapes of transcriptome, epigenome, and 3D genome in mouse brain single cells

Cell-type specific transcriptional programs are pivotal in determining cell lineage and delineating the intricate relationship of cell identity and functions within complex tissues or organs. In the studies of mouse brain, diverse cell populations based on gene expression have been extensively documented(*34–39*), establishing a rich knowledge base. However, the genomic mechanisms that modulate cell-specific transcriptional programs are still vague. In this direction, we positioned ourselves to study the intricate interplays between gene expression, epigenomic state and chromatin folding structure. To achieve this, we harnessed the high-throughput capability of ChAIR and conducted a comprehensive profiling of the 3D epigenomic (3D chromatin folding and epigenomic states) landscape and transcription regulation in mouse brain cells. We stratified our study by selecting five critical developmental age points from infancy through young adult to aged mice: postnatal days 2 (P2), 11 (P11), 95 (P95), 365 (P365, 1-year), and 730 (P730, 2-year) (Fig. 2A), aiming to gain a thorough understanding how 3D epigenomic states may coordinate and influence transcriptional changes during developmental and aging processes in the brain.

**Fig. 2.**
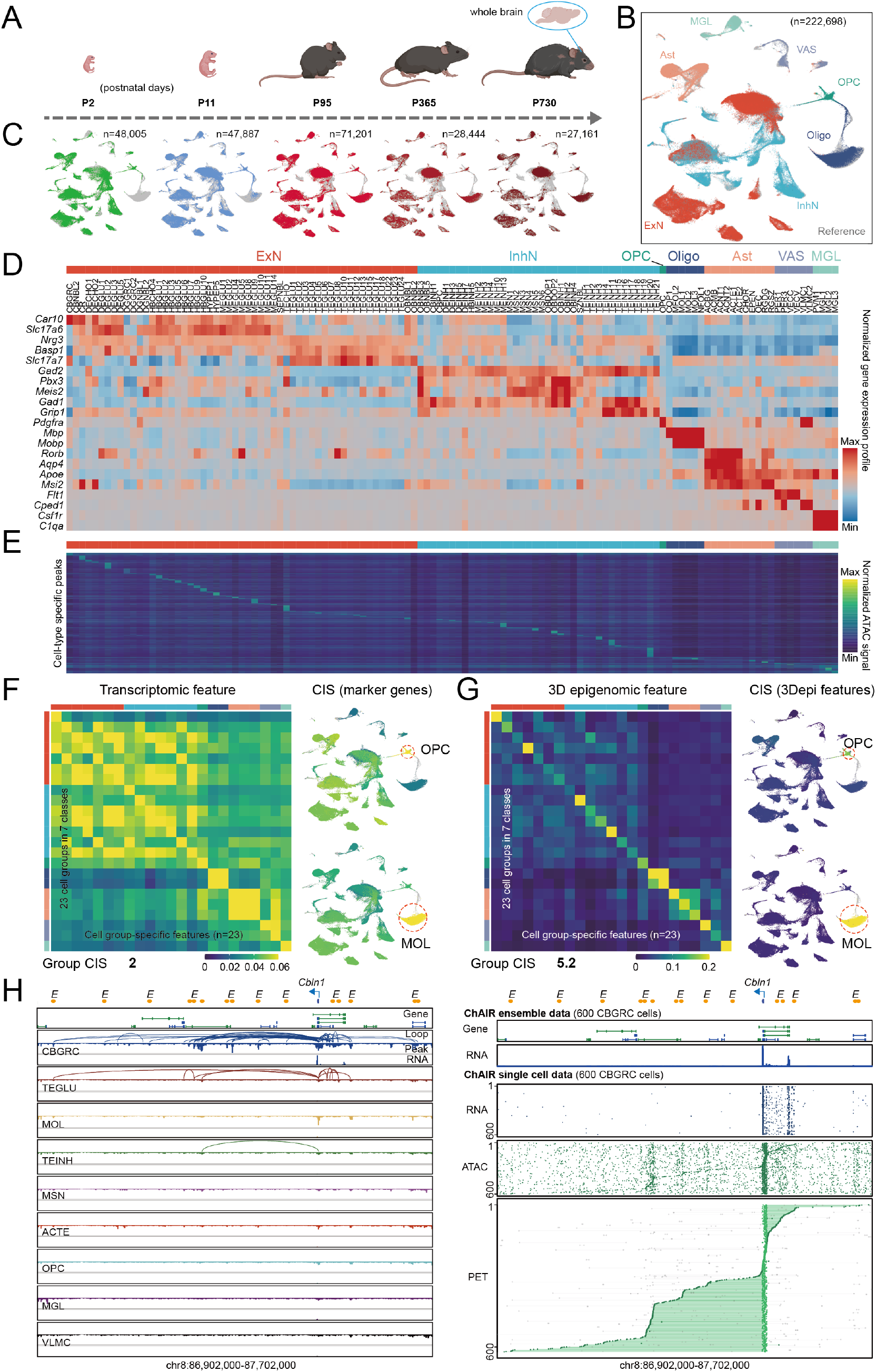
Single-cell 3D epigenome and transcriptome landscapes of whole brain cells during mouse maturation and aging. **(A)** The study design of collecting whole brain tissues at various postnatal days. (**B**) UMAP plot of combined ChAIR-RNA data highlighting major brain cell classes in various colors and reference in grey. The number (n) of cells was provided. (**C**) UMAP plots of ChAIR-RNA data in 5 age points. The cell numbers (n) were provided. **(D)** Matrix plot of canonical marker gene expression profile in 121 brain cell types of 7 cellular classes. (**E**) Matrix plot of cell-type-specific ChAIR-ATAC peak signals across 121 cell types. **(F-G)** Matrices of cell identity score (CIS) calculated using transcriptomic (**F**) and 3D epigenomic features (**G**) across 23 cell groups in ChAIR data along with UMAP plots showing the specificity revealed by transcriptomic feature (**F**) and 3D epigenomic feature (**G**) for OPC and MOL cells. (**H**) Browser view of ensemble ChAIR data for CBGRC-specific enhancers (left) along with the single-cell ChAIR data tracks at the *Cbln1* locus (right).

With rigorous quality control measures (see methods), we collected high-quality ChAIR data of 222,698 cells sampled from the 5 age points with high reproducibility (table S1, fig. S10). We annotated the ChAIR-RNA data using a label transfer method(*40*) based on a reference scRNA-seq dataset(*34*) that profiled central nervous system cells of mouse brain. The combined ChAIR-RNA data showed a high degree of similarity with the reference data (Fig. 2B and fig. S11A), while distinct age-related variations were discernible across different age points (Fig. 2C and fig. S10A). Of the 199 well-defined mouse brain cell types delineated in the reference(*34*), we identified 121 cell types characterized by the expression of canonical marker genes (Fig. 2D and fig. S11B), which belong to 45 cellular groups in 7 major brain cell classes: excitatory neurons (ExN), inhibitory neuron (InhN), astroependymal (Ast), oligodendrocyte progenitor cells (OPC), oligodendrocyte (Oligo), microglia (MGL), and vascular cells (VAS) (Fig. 2D and fig. S11A). Meanwhile, each cell type identified through ChAIR-RNA data featured corresponding high-quality ChAIR-ATAC and -PET datasets. These datasets demonstrated a high correlation of epigenomic state and chromatin folding features with related methods across major cell classes(*39, 41*) (fig. S12A-B and fig. S13A), and showcased specific open chromatin loci (Fig. 2E and fig. S12C) and chromatin folding features in compartment, TAD, and chromatin contacts in ensemble and single-cell views (fig. S13B-C).

To analyze the correlation of RNA, ATAC, and PET data generated by ChAIR, we focused on 23 out of the 45 cell groups, in which each of the trimodal datasets were sufficient for in-depth analysis. With a focus on cell-type specific marker genes, we reasoned that the 3D epigenomic features in individual cells should exhibit a degree of specificity analogous to that observed in RNA for defining cell identity. To quantitatively evaluate each data modality for cell type identification, we developed a scheme to calculate cell identity score (CIS) that measures the distinctiveness of a given feature, including transcription (RNA data), gene activity (ATAC signal on gene body), promoter strength (ATAC at TSS), enhancer strength (ATAC at enhancer site), and chromatin loop (PET data), in a specific cell type over the background noise in other cells (see methods). All mono-modal CIS displayed observable cell-type specificity, with the transcription CIS and chromatin loop CIS values were at the same level, whereas ATAC-related CISs lagged behind (fig. S13D). It is noteworthy that although the CIS value for the specificity of transcription and the chromatin loop showed no significant difference, their matrix profiles exhibited distinct differences, suggesting that chromatin contacts may provide a perspective for cell identification that is different from gene expression. Furthermore, integrating chromatin loop specificity with ATAC-based open chromatin specificity, collectively referred to as 3D epigenomic features or states, significantly enhanced the CIS value for each specific cellular group and all 23 cellular groups collectively (see methods) (Fig. 2F-G and fig. S13D). For instance, in the case of OPC and mature oligodendrocytes (MOL), the composite CIS of the 3D epigenomic feature were substantially higher, providing a clearer distinction between specific signals and background noise, than those derived solely from transcriptomic feature for both OPC and MOL, respectively (Fig. 2F-G).

The 3D epigenomic landscape associated with cell specific marker genes enabled us to identify cell-type specific enhancers that may have practical potential to optimize the manipulation of exclusive transcription activation in functional neuroscience and therapeutic development(*42–44*). We examined the likely cis-regulatory elements (CREs) associated with the marker genes in each of the cell groups (n=23) with a set of stringent criteria (see methods). This approach led to the identification of 562 high-quality distal enhancer loci that displayed high cell-type specificity (table S2 and fig. S14A-J). A unique example involved 12 notable enhancers specifically connected to the promoter of *Cbln1* (Fig. 2H). This gene encodes a protein Cerebellin-1 essential for synaptic integrity and plasticity specifically in cerebellar granule cells(*45*). Interestingly, while the *Cbln1*-associated TSS-E loops were also observed to some extent in TEGLU, a different type of granule cell in the cerebral cortex, they were not sufficient to activate *Cbln1* expression (Fig. 2H, left), further highlighting the specificity. The single-cell-resolved browser views of ChAIR data at the *Cbln1* locus further revealed extensively concerted chromatin interactions, high level of open accessibility, and active transcription (Fig. 2H, right). More examples of cell-type specific TSS-E interactions were presented in fig. S14K.

Thus, our observations demonstrated a strong link between genomic structure and function, crucial for defining cell identity. Importantly, marker gene-associated 3D epigenomic states robustly indicate cell identities, emphasizing that the combined epigenomic states and 3D chromatin contacts are closely linked with cell-specific transcriptional programs.

### Spatially-resolved regional specificity of 3D-epigenomic features

ChAIR is efficient in generating tri-modality data including RNA and genomic information; however, it does not offer anatomical information, unlike spatial transcriptomics. While efforts had been made in combining the robustness of single-cell techniques with the positional details of spatial analysis (*46*), mapping cell-type specific genome conformations within a spatial framework still faces considerable challenges. To address this, we utilized a spatial transcriptomic dataset from a mouse coronal hemibrain section generated by Stereo-seq from an adult mouse(*47*). Our goal was to incorporate the 3D epigenomic features of the ChAIR data with the brain regional specificity provided by the Stereo-seq data. We reasoned that if the ChAIR-RNA data matched with the Stereo-seq RNA data, then the concurrent ChAIR-ATAC and -PET data could be assigned to the corresponding spatial positions defined by Stereo-seq, thereby adding a crucial spatial dimension to our understanding of 3D epigenomics.

We first annotated the bin 50 (50 × 50 bins) data in Stereo-seq using the ChAIR-RNA data (see methods) and successfully linked 20 anatomical regions identified by Stereo-seq to 19 cell types defined by ChAIR-RNA data with high specificity (Fig. 3A and fig. S15A-C). These cell types, especially neurons from cortex and hippocampus, closely matched with their original spatial annotations(*34*). We validated the ChAIR-Stereo integration by assessing the spatial distribution of canonical marker genes in both Stereo-seq data and the histological staining of adjacent tissue section from Allen Brain Atlas (fig. S15D). Next, we mapped the ChAIR-RNA signals of marker genes in various ExN cells, including TEGLU and DGGRC, onto the anatomical profile of a coronal hemibrain section in Stereo-seq (Fig. 3B). These results demonstrated unique expression patterns across the cortical layers, highlighting the distinct regional characteristics of ExNs in outer (layer 2/3), middle (layer 4), and inner (layer 5/6) regions of neocortex, as well as in the dentate gyrus of hippocampus (Fig. 3B, 1-2^nd^ columns), indicating high accuracy of ChAIR-Stereo data integration.

**Fig. 3.**
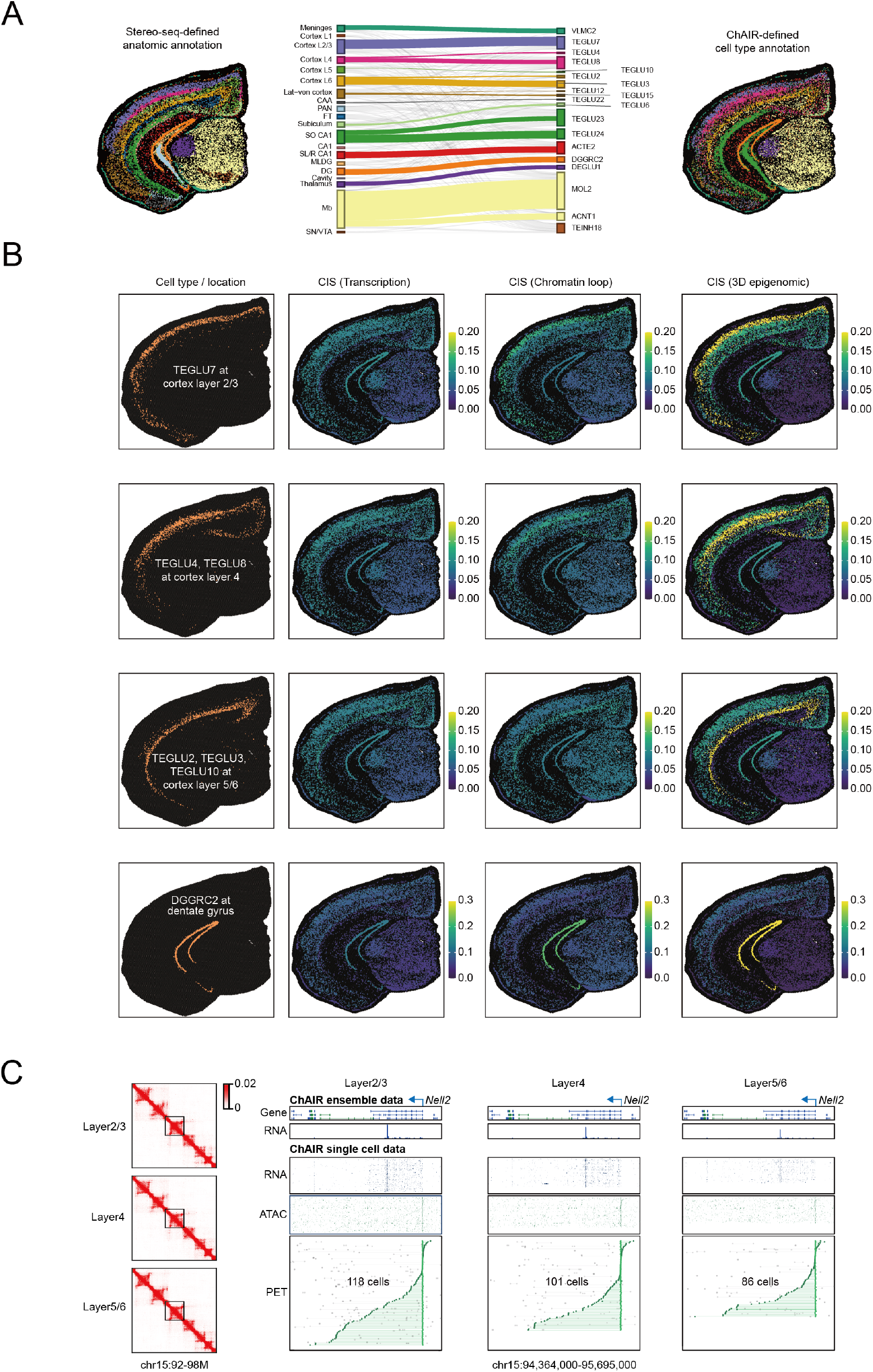
Regional-specific 3D epigenomic features in mouse brain cells revealed by ChAIR. (**A**) Integration of Stereo-seq (bin50) data with ChAIR-RNA data, connecting regions in a coronal hemibrain section defined by Stereo-seq (left) with the corresponding cell types characterized by ChAIR (right). (**B**) Spatial distribution of specific cell types and locations (1^st^ column) and the corresponding signal intensity of CIS score calculated by marker gene transcription (2^nd^ column), marker gene associated chromatin loop (3^rd^ column), and 3D epigenomic (3Depi) (4^th^ column) features derived from ChAIR data. (**C**) An example of ChAIR data showing 2D contact maps and the single-cell tracks of RNA, ATAC, and PET at the locus *Nel2* (a cortex layer 2/3 marker gene) in layer 2/3, layer 4, and layer 5/6.

In characterizing the ChAIR data, we had realized that chromatin loops are also distinctive features, and the composite 3D epigenomic features provided a superior specificity for cell identity (Fig. 2F-G and fig. S13D). Specifically for the ChAIR data that were matched the Stereo-seq data, we focused on 8 groups of cells that have sufficient data 3D epigenomic data for in-depth analysis (fig. S15E). Strikingly, the chromatin loop and specifically the 3D epigenomic features exhibited significantly higher CIS values for than those represented by gene expression alone (Fig. 3B, 2-4^th^ columns, fig. S15E, and S16). For example, the marker gene transcription profile of TEGLU7 (specific in cortex layer 2/3) showed signal enrichment in both cortex layer 2/3 and dentate gyrus, while the chromatin loop and the 3D epigenomic features exhibited particularly higher signals in cortex layer 2/3. Similarly, while the marker gene transcription of DGGRC cells showed enriched signals at hippocampus region, it also showed high-level of gene expression in cortical layers. However, the chromatin loops and 3D epigenomic profiles were highly specific to the hippocampus with minimal background noise in other anatomic regions. Lastly, the single-cell-resolved browser views, notably at the *Nell2* locus—a marker gene for cortex layer 2/3 (Fig. 3C) and additional examples (fig. S17)—provided detailed insights, highlighting specific marker genes with associated chromatin interaction loops and ATAC signals.

Thus far, our results demonstrated a technical advance that RNA-based cellular specificity could be better defined by the integrated 3D epigenomic features and be accurately mapped onto spatially-resolved anatomic regions using ChAIR-RNA data as a bridge. Additionally, the better performance of 3D epigenomic profile in defining cell identity further suggested that 3D epigenomic state is closely linked with gene transcription(*19*).

### Chromatin rewiring and associated transcriptional changes during aging

From the annotated ChAIR-RNA data, we explored the dynamic shifts of cellular composition within the whole mouse brain across lifespan and observed a notably higher proportion of neuroblasts (progenitors of excitatory and inhibitory neurons) early in life (P2-P11), and nearly disappeared after maturation (Fig. 4A), highlighting an intense period of early neural development. The proportion of mature neurons also showed substantial changes, with the ExNs increasing and the InhNs decreasing proportionally over time. For non-neuronal cells, there were notable changes as well, particularly the balance between OPC and oligodendrocyte. Interestingly, the proportion of microglia increased in aged brains, suggesting an upregulation of immune surveillance or neuroinflammatory responses as mouse ages(*35*), whereas the quantities of vascular cells and astrocytes remained remarkably constant throughout the lifespan, underscoring their critical and unchanging roles in maintaining neural health and functionality (Fig. 4A). Furthermore, the dynamic shift in cell composition also occurred in different brain regions. The most pronounced change was seen in the distribution of ExNs in telencephalon (TE) and cerebellum (CB). In early infant time (P2), the majority of ExNs (60%) were found in TE and only 10% in CB, then quickly shifted to 35% vs 50% in P11, and the cerebellar granule cells (CBGRC) gradually became the predominant ExNs (more than 80%) in later life (Fig. 4B, left). Conversely, the proportion of InhNs remained relatively stable, showing no significant changes (Fig. 4B, right). These observations based on ChAIR-RNA data revealed a complex dynamic among different cell types over the course of brain maturation and aging, highlighting substantial changes in transcriptional programs that define cellular properties. This presents an ideal opportunity to investigate the underlying genomic mechanisms using the multi-omic ChAIR-ATAC and -PET data captured in the same cells.

**Fig. 4.**
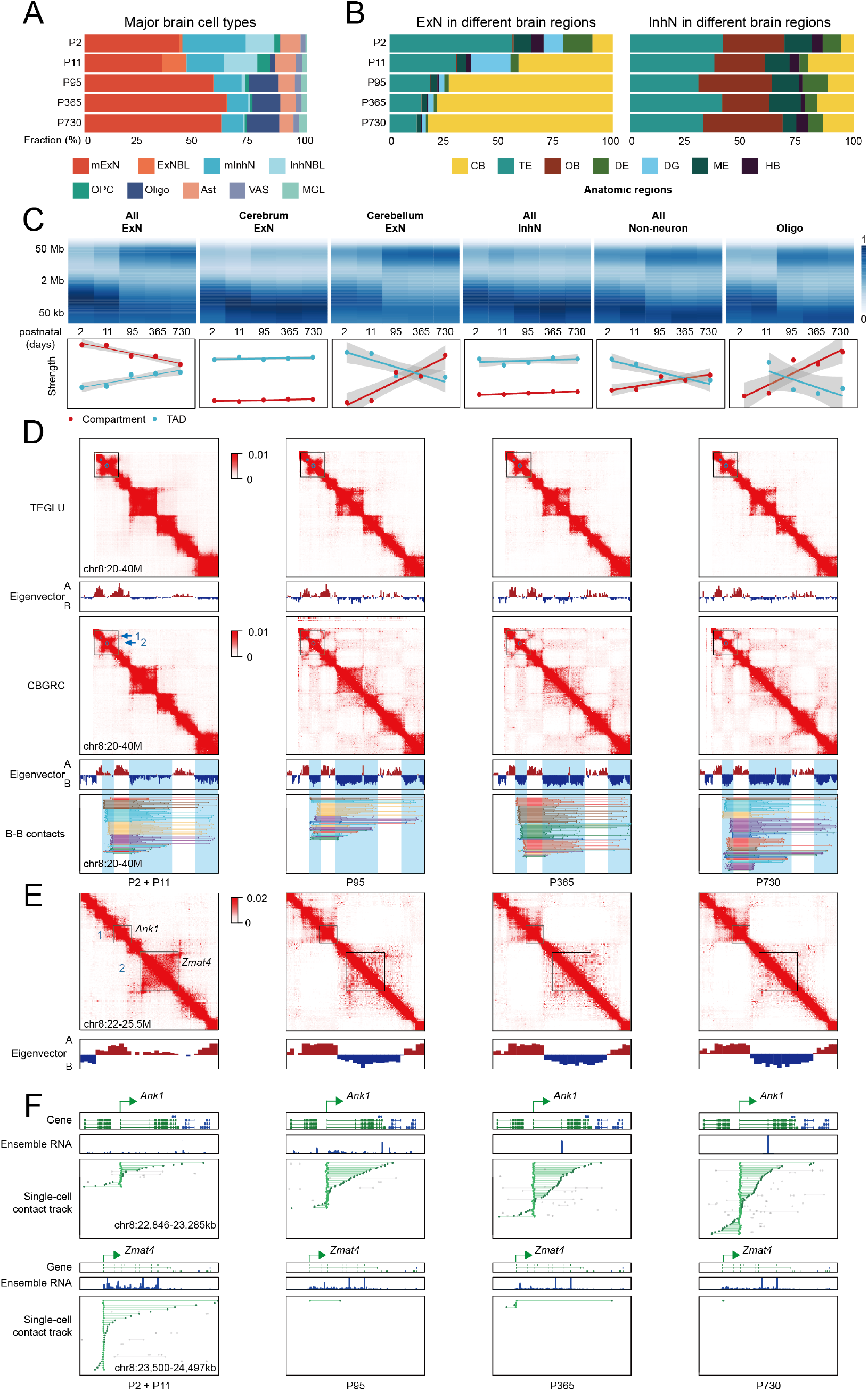
Aging-related remodeling of chromatin folding in mouse brain cells. (**A**) The dynamic shifting of cellular composition of major brain cell classes and their progenitors during maturation and aging from infant (P2, P11) to adult (P95) and older stages (P365 and P730). mExN, mature excitatory neurons; ExNBL, excitatory neuroblast; mInhN, mature inhibitory neurons; InhNBL, inhibitory neuroblast. (**B**) The dynamic changes of neuronal composition in different brain regions across 5 age points. CB, Cerebellum; TE, Telencephalon; OB, Olfactory bulb; DE, Diencephalon; DG, Dentate gyrus; HB, Hindbrain (Pons and Medulla); ME, Mesencephalon. **(C)** The distance spectrum of chromatin contacts in different cells across 5 age points (top) from infant (P2, P11) to adult (P95) and older stages (P365 and P730) along with aggregated strength of A/B compartment and TAD signals (bottom). (**D**) Example views of ChAIR-PET data in 2D contact heatmaps (chr8:20-40Mb) and Eigenvector for compartments A/B in TEGLU and CBGRC cells, focusing on the *Ank1*(1) and *Zmat4*(2) gene loci in the segment of chr8:20-40Mb across age points (top). The single-cell browser views of ChAIR-PET data, showing contacts between B compartments were also shown (bottom). (**E**) Zoom-in views of 2D contact heatmaps from the boxed area in (D), showing a compartment switch from A to B and the dissolution of the TAD structure around *Zmat4* during aging. (**F**) Browser views of single-cell chromatin contact tracks of ChAIR-PET data in CBGRC cells at the *Ank1* (top) and *Zmat4* (bottom) loci across age points.

Previous findings suggested that the spectrum of chromatin contact distance is a characteristic measurement of genome conformation in different brain cells(*48*). Using ChAIR-PET data, we examined the distribution of chromatin contact distances in various types of mouse brain cells and revealed distinct patterns over the 5 age points: ExNs and non-neuronal cells exhibited predominantly short-range contacts (20kb-2Mb) in infant time (P2/11). Notably, as these cells matured and aged (P95-730), there was a significant shift toward predominantly ultra-long chromatin contacts (>2Mb, hereafter referred to as ‘megacontacts’), whereas InhNs consistently maintained short-range contacts over time (Fig. 4C and fig. S18A-B). Further dissection of the data pinpointed that the change of chromatin contact ranges in ExNs was specific to cerebellum, while cerebrum remained constant over time. This observation aligns with a recent finding that cerebellar granule cells undergo gradual chromatin remodeling throughout their lifespan, eventually exhibiting predominantly ultra-long megacontacts in aged cells(*49*). However, for non-neuronal populations, only Oligos displayed a notable pattern, shifting markedly from short-range in early life to predominance by megacontacts in mature and aged mice (Fig. 4C). Intriguingly, chromatin conformation in other non-neuronal cells remained relatively constant over time either as short contacts in OPCs or as megacontacts in VAS and MGL cells (fig. S18A-C). Additional analyses revealed that cells with chromatin contacts predominantly in short-range were positively correlated with RNA expression and negatively with trans-contacts. Conversely, cells with extensive megacontacts demonstrated a negative correlation with gene expression and positive with higher frequency of trans-contacts (fig. S18D). These observations suggested that cells with dominant short-range of chromatin contacts are in a more transcription-active chromatin state than those with ultra-long megacontacts.

To characterize chromatin’s conformational change in details, we examined chromatin folding structures at various scales, including compartments, TADs, and loops using ensemble and single-cell ChAIR-PET data (Fig. 4D-F and fig. S19-20). Our findings indicated that short-range chromatin contacts (20kb-1Mb) typically correlated with a predominance of TAD structures, while megacontacts were associated with increased compartmentalization in brain cells as they age (Fig. 4C, bottom and fig. S19). More specifically, the chromatin conformation in ExNs from cerebrum or telencephalon (TE) displayed a consistent level of TAD and compartment signals, whereas the cerebellar ExNs were characterized by a notable transition from a TAD-dominant state to a state with increased compartmentalization over time (Fig. 4C, bottom and fig. S19). A 20 Mb segment on chr8 exemplified the characteristics of different chromatin structures in TE and CB. The 2D contact heatmaps revealed stable, well-defined TAD structures in TEGLU cells over time (Fig. 4D top). In contrast, CBGRC cells showed a marked shift from TAD predominance in infancy (P2/11) to a compartment-dominant state with distinct plaid patterns in adulthood (P95) and old age (P365/730) (Fig. 4D top). This increased compartmentalization was highlighted in browser views with single-cell resolution (Fig. 4D, bottom and fig. S18E).

Further analyses of a zoomed-in 500 kb genomic region featuring *Ank1*, which encodes a protein enriched in the cerebellum(*50*) and a gene *Zmat4,* known for its role in Alzheimer’s disease(*51, 52*). The chromatin where *Ank1* resides maintained in active state, and the associated TAD structure appeared to expand with distinguishable stripes after maturation, especially in aged CBGRC cells (Fig. 4E). Concurrently, *Ank1*’s expression and the number of chromatin loops linked to its promoter consistently increased over time (Fig. 4F, top). Interestingly, the nearby chromatin domain that hosts *Zmat4* underwent a notable compartment switch from A in early life (P2/11) to B in later stages. This transition was also accompanied with the dissolution of the TAD structure (Fig. 4E) and the disappearance of promoter-associated chromatin interactions (Fig. 4F, bottom). These conformational changes likely served as the structural basis for high level expression of *Zmat4* in early life and decreased in later stages (Fig. 4F, bottom). Similarly, on chr18, the *Dcc* gene, which encodes netrin-1 receptor crucial for axon guidance(*53*), also underwent a compartment switch during aging, leading to increased B-to-B compartment contacts (fig. S20A) and reduced TAD features (fig. S20B). The single-cell browser views further delineated robust ChAIR signals for *Dcc*-associated RNA, ATAC, and chromatin interactions in early life (P2/11), which diminished in old age (fig. S20C).

Together, our findings suggest that the transition from TAD-dominant structures with short chromatin contacts in early life to compartment-dominant structures with ultra-long megacontacts in later life serves as a genomic indicator of cellular aging. Additionally, the age-related chromatin features closely coincided with transcriptional activity, suggesting a likely causal relationship between the chromatin structure and the regulation of aging-related genes. Importantly, our analyses revealed a complex pattern of aging in ExNs across different brain regions. Specifically, the limited flexibility in chromatin conformational changes observed in TEGLU may indicate reduced adaptability to the aging process. In contrast, the extensive chromatin conformation shifts in the CB could provide a genomic mechanism that condenses chromatin structure, potentially leading to reduced nuclear size and corresponding transcriptional changes in aging-specific genes.

### Concerted dynamics of chromatin folding and transcription during brain cell differentiation

It has been shown that mouse brain cells undergo substantial changes in transcription and genome architecture during neuronal development(*41*). However, most of the prior knowledge had been derived from partial brain tissues and separated experiments followed by integration of mono-modality datasets with many prior assumptions. Here, we aimed to investigate the relationship of transcriptome, chromatin accessibility, and 3D genome structure across different lineages using the tri-omic ChAIR data derived from whole mouse brains.

We first surveyed the well-documented cellular lineage from OPCs to mature oligodendrocytes(*34*). All OPCs and oligodendrocyte cells were ordered according to the proportion of megacontacts from the least to the most in individual cells (Fig. 5A). This ordering effectively recapitulated the differentiation trajectory from OPC to mature oligodendrocyte, revealing a striking concordance between RNA pseudotime and the sequence determined by chromatin conformation (hereafter referred to as ‘chromatin contact pseudotime’) (Fig. 5A-B). In this arrangement, OPC predominantly featured short-range contacts at the lower end of the distance spectrum (Fig. 5A, bottom), while most MOL1 cells exhibited megacontacts in stark contrast. To confirm these bi-polar trends, we assessed the ChAIR data for expression (RNA), chromatin accessibility (ATAC), and promoter-linked chromatin contacts (PET) associated with the marker genes in OPC and MOL1 cells (Fig. 5B-C), respectively. The results showed that the marker gene associated features correlated well with how the cells were arranged based on chromatin contact pseudotime (Fig. 5C and fig. S21A). Furthermore, our 3D modeling results (see methods) indicated that the nucleus of OPC is significantly larger than MOL1 (Fig. 5D, fig. S21C).

**Fig. 5.**
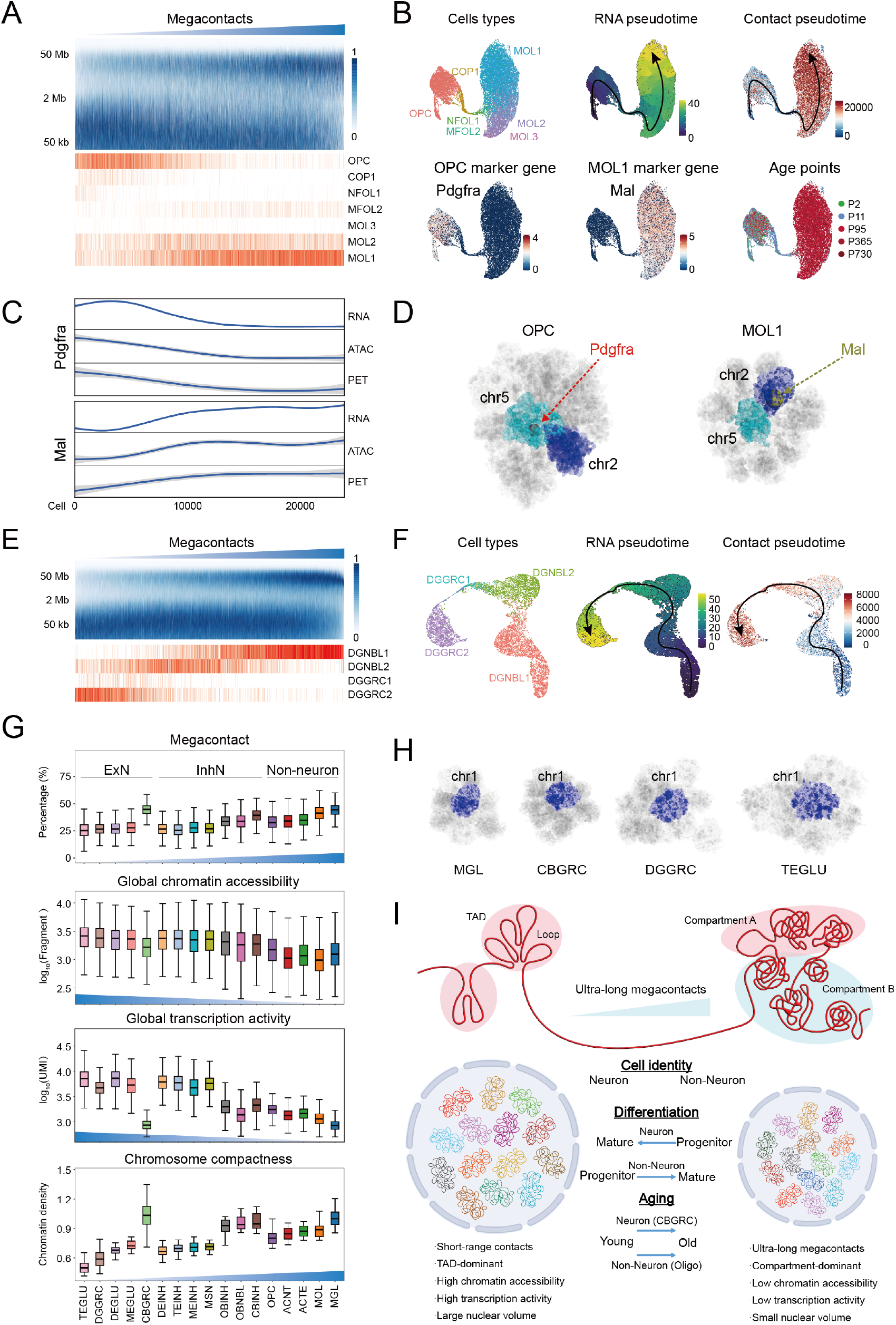
Chromatin architecture reorganization during cell differentiation. (**A**) The spectrum of chromatin contact distance of OPCs and oligodendrocytes. Individual cells (vertical lines, n=23,904) were sorted by the extent of megacontacts from the least to the most (top). OPC and subtypes of oligodendrocytes in coordination with the sorting of chromatin contact distances (bottom). (**B**) UMAP plots of ChAIR data by several analyses (from left to right): cellular lineage from OPCs to oligodendrocytes, RNA pseudotime, chromatin contact pseudotime, OPC-specific marker gene *Pdgfra* profile, MOL1-specific marker gene *Mal* profile, and cell distribution of age points. (**C**) Correlation of chromatin contact distances with marker gene expression (RNA), promoter-associated chromatin accessibility (ATAC), and promoter-associated chromatin interactions (PET) in individual cells. (**D**) The example views of reconstructed 3D genome models of OPC and MOL1 metacells. Nuclear positions of specific marker genes and chromosomes were indicated. The sizes of balls reflected the relative levels of gene expression. (**E**) The spectrum of chromatin contact distance (top) and cellular composition (bottom) of neuroblast DGNBL1/2 to DGGRC1/2 (n=8417) in dentate gyrus. (**F**) UMAP plots of ChAIR data (from left to right) on the differentiation trajectory from DGNBL1/2 to DGGRC1/2, RNA pseudotime, and chromatin contact pseudotime. (**G**) Boxplots (from top to bottom) of the megacontact percentage, global chromatin accessibility, global transcription activity, and chromosome compactness (chromatin folding density) of 17 specific cell types ExNs, InhNs, and non-neurons. (**H**) The examples of reconstructed 3D architectures representing nuclei among various brain cells, with microglia (MGL) having the highest density and smallest volume and TEGLU cells having the lowest density and largest volume. (**I**) A generalized working model depicting the role of chromatin folding profiles with short-range contacts and ultra-long megacontacts in different mouse brain cells and during various biological processes.

We then extended our analysis to examine the cellular differentiation lineage of ExNs in dentate gyrus (DG), tracing the progression from neuroblasts (DGNBL1/2) to mature granule cells (DGGRC1/2)(*54*). Intriguingly, the chromatin contacts in most DGNBL1 cells were predominantly in ultra-long range, while DGNBL2 positioned in the middle, and DGGRC1/2 cells exhibited mostly short-range contacts (Fig. 5E, bottom). This genomic trajectory from DGNBL to DGGRC stood in contrast to the pattern observed in the differentiation of OPC to MOL. To validate the genomic trajectory observed from DGNBL to DGGRC, we examined RNA, ATAC, and PET profiles associated with the marker genes specific to DGNBL and DGGRC(*54*) (table S3, fig. S21B, and S22A) and the estimated nuclear sizes (fig. S21C, and S22B). These analyses confirmed the authenticity of our observations, suggesting an intriguing notion that neurons and non-neuronal cells in the mouse brain may undergo different genomic mechanisms during their differentiation.

To substantiate our observations of ExNs in DG, we analyzed additional neuronal differentiation processes from cerebellar neuroblasts (CBNBL2) to granule cells (CBGRC) in CB. The ordering of CBGRC cells (P2-P730) based on chromatin contact aligned with the patterns we observed during aging (Fig. 4C) and revealed that CBNBL2 differentiated to CBGRC early in life (fig. S22C-F). Focusing on the infant (P2) ChAIR data, we noted that chromatin contacts in CBNBL2 predominantly fell within the ultra-long range as megacontacts, while CBGRC cells were primarily associated with short chromatin contacts (fig. S22C). Additionally, the trajectories of RNA pseudotime, chromatin contact pseudotime, and marker gene expression profiles were closely aligned (fig. S22D) and matched with the patterns observed in DG (Fig. 5E-F). Similar patterns were also observed in InhNs from olfactory bulb (OB), specifically between OBNBL3 and OBINH2 (fig. S22G-J). Thus, neurons located in different brain regions displayed the same patterns during differentiation, progressing from progenitor cells to mature neurons with accompanying chromatin remodeling from megacontacts to short contacts, further confirming the notion that neurons possess different genomic properties and behaved differently than non-neuronal cells.

Recognizing the potential implication of megacontacts in deciphering the mechanisms of differentiation and aging (Fig.4 and Fig.5A-F), we sought to assess their distribution in different types of post-mitotic mouse brain cells. In adult (P95) mouse brain, our ChAIR data showed predominant megacontacts in the genomes of non-neuronal cells, whereas the neurons were enriched with short-range contacts, except CBGRC as an outlier (Fig. 5G). A clear trend was observed that the short-range contacts were associated with higher global chromatin accessibility and transcription activity, and lower chromatin folding density. In contrast, a higher megacontact ratio was linked to decreased global chromatin accessibility and transcription activity (Fig. 5G), consistent with what had been hinted in previous mapping-based studies(*41, 48, 55*). Our observation suggests that ultra-long megacontacts may indicate to an overall genome compaction and transcription reduction. Importantly, the chromatin density estimated from our 3D modeling analysis, based on ChAIR-PET data, closely correlated with the ratio of megacontacts in a genome (Fig. 5G-H, fig. S23). This implies that neurons, particularly excitatory neurons, have lower genome folding density and larger nuclear volume than non-neuronal cells. Coincidentally, our findings based on mapping also aligned with a recent observation in mouse brain cells based on an imaging method DNA-MERFISH (*56*). Among other non-neuronal cells, MGLs displayed the smallest nuclear volumes, whereas InhNs and ExNs exhibited significantly larger volumes. Thus, our analyses underscored that the global chromatin contact profile, particularly the megacontact ratio in a genome, is a crucial metric that reflects not only the genome compaction and nuclear size but also the overall transcriptional state of chromatin and distinct cellular identities, profoundly influencing essential biological processes such as differentiation and aging.

## Discussion

In this study, we introduced a single-cell method (ChAIR) that simultaneously maps chromatin accessibility, interaction, and RNA expression in individual cells. This innovative approach provided several technical advances over the current multi-omic single-cell methods in several aspects. Existing methods like ISSAAC-seq(*24*) and SHARE-seq(*25*) capture RNA and ATAC together but lack chromatin contact information, preventing them from directly linking distal elements in open chromatin loci to transcription regulation. Meanwhile, methods such as HIRES(*19*) and scCARE-seq(*18*) detect RNA and Hi-C together but fall short in providing epigenomic information for accurately identifying genomic elements that are specifically involved in transcriptional programs. The obvious advantages of ChAIR by capturing RNA, ATAC, and PET data concomitantly within the same cells is its ability to identify active genes, concurrent open chromatin states of regulatory elements, and chromatin interactions between gene promoter and distal regulatory elements, thus providing an integrated 3D epigenomic state to elucidate the underline genomic mechanism for gene transcription regulation.

Another significant advantage of ChAIR over existing methods is its microfluidic-based high-throughput capability. A single ChAIR experiment, utilizing multiple microfluidic channels, can efficiently generate up to 100,000 high-quality single-cell datasets. This feature is particularly advantageous for analyzing complex tissues and organs, such as brains and tumors, which contain a wide array of diverse cell types across various developmental stages, aging processes, and environmental conditions. Therefore, the high-throughput capacity of ChAIR is not only enabling the comprehensive profiling of cellular heterogeneity but also unraveling the intricate interplays between different but highly related cells within a microenvironment of a complex biological systems and the underline regulatory mechanisms.

Lastly, ChAIR data can be readily integrated with spatial transcriptome data, providing spatially-resolved specificity. Recent advances in spatial transcriptomics allow for *in situ* mapping of gene expression within the tissue context, adding a new dimension to single-cell biology. While current spatial methods are effective for transcriptome mapping, they typically lack the capability to provide a comprehensive spatial genomic map with high resolution. By integrating ChAIR data with spatial transcriptomic data derived from the same tissue samples, we can directly align the ChAIR-RNA data with the spatial RNA data. This convenience enables the precise positioning of ChAIR-identified cells, along with their concurrent 3D epigenomic structures, within specific anatomical regions on the spatially-resolved transcriptome map. This integration not only enhances the specificity of spatial genomic analyses but also facilitates deeper mechanistic studies of transcription regulation by correlating 3D epigenomic profiles with spatial tissue contexts.

A primary limitation of the current ChAIR, as with all multi-omic methods, is its relatively lower efficiency compared to mono-omic techniques. While the ChAIR-RNA data matches the quantity with other methods, the yield of DNA data tends to be lower. This is partly due to the inherent characteristics of high copy numbers of transcripts from active genes versus only two copies of DNA templates. Consequently, using ChAIR-ATAC and ChAIR-PET data for unsupervised cell clustering analysis is generally less effective than using RNA data alone. However, the high-throughput nature of ChAIR would compensate this issue. Based on ChAIR-RNA clustering, cells with closely related gene expression profiles were brought together into metacells. Usually, a metacell with a few hundreds of single cells would gather sufficient ChAIR-ATAC and ChAIR-PET data enough for genomic analysis and meanwhile avoid individual stochasticity. Particularly, by leveraging marker genes specific to each of the metacells, the composite 3D epigenomic features—combining marker gene promoter-associated chromatin interactions and ATAC signals—demonstrated significantly higher specificity and reduced non-specific background noise compared to using RNA data alone in defining cell identity. Ongoing refinements to the ChAIR protocol and data processing pipeline will render ChAIR improved efficiency and enable deeper inquiring for mechanistic insights into the genomic underpinnings of gene transcription regulation.

To explore the genomic dynamics of mouse brain cells during maturation and aging, we applied ChAIR to whole brains of mice ranging from postnatal 2 days to 2 years old. Our comprehensive ChAIR data captured the progressing single-cell landscapes of the 3D epigenome and transcriptome across the lifespan. Notably, the integrated analyses of concurrent RNA, ATAC, and PET data from individual cells revealed tightly concerted patterns of epigenomic states, 3D chromatin conformation, and gene expression in various cellular lineages and in the process of growing old, aging. This close coordination in single cells strongly suggest a possible causal relationship between chromatin folding states as the structural basis and the transcription as a functional output. However, the true nature of these coupled events will have to be uncovered by functional perturbation assays in single cell and in single molecule levels.

Specifically, we observed that ultra-long megacontacts (>2Mb) generally correlate with aging, as demonstrated in this study and partly supported by other reports(*49, 57*). Particularly, the genomes of early-life brain cells are characterized by predominantly short-range contacts, which gradually shift to ultra-long-range contacts later in life. This trend is also observed during cellular differentiation of non-neuronal cells from progenitors to mature cells, such as the transition from OPCs to mature oligodendrocytes (Fig. 5A). However, in contrast, neuroblasts initially displayed more megacontacts but transitioned to became primarily short-range contacts when they mature (Fig. 5E-F and fig. S22) as we demonstrated for neurons in brain regions of dentate gyrus, cerebellum, and olfactory bulb.

Further analyses of genomes with varying chromatin contact distances across different cell types revealed distinct patterns: genomes with short contacts are TAD-dominant, featuring more open chromatin and higher transcriptional activity, while those with a higher proportion of megacontacts tend to be more compartmentalized with condensed chromatin state and reduced gene expression. Therefore, the presence of megacontacts serves as a reliable indicator of the overall 3D epigenome states and associated cellular properties during differentiation, across various cell types, and throughout the aging process.

Additionally, using an established 3D modeling tool(*14*), despite the absence of haplotype-resolved data, we reconstructed the 3D genome structures, calculated the folding density of individual chromosomes, and estimated nuclear volume for various cell types and at different age points. We observed a strong correlation where a higher proportion of megacontacts in a genome is associated with increased genome density, or inversely, with reduced nuclear volume. This finding aligns with the estimations of nuclear sizes in various brain cell types by a recent imaging-based analysis(*56*). Thus, global chromatin contact distance not only serves as an effective indicator of genome conformation, correlating with detailed 3D epigenomic features such as chromatin folding structures and transcription activities, but also acts as a convenient surrogate for assessing nuclear morphology in individual cells (Fig. 5I).

It is noteworthy that the dynamic balance between TEGLU in TE and CBGRC in CB proportionally across five age points (Fig. 4B) may be influenced by a mechano-genomic mechanism. In comparison, the genomic properties between TEGLU and CBGRC are in sharp contrast. TEGLU maintained shorter chromatin contacts and higher transcription activity along with lower chromatin folding density or larger nuclear size, while CBDRC exhibited exceptionally high proportion of megacontacts and high chromatin density along with extremely low transcription (Fig. 5G, fig. S23). A plausible explanation for the dramatic shift in cellular composition between ExNs in TE and CB (Fig. 4B) is that the inflexible chromatin conformation and larger nuclear size may make TEGLU more susceptible to apoptosis during course of aging. Conversely, the gradual increase in megacontacts, genome compaction, and the reduction of transcription activity and nuclear size (fig. S23) may render CBGRC better adapted in the process of senescence for energy conservation(*49*), thus enhancing survival as the mouse ages. Functionally, this observation aligns with the evolutionary perspective, as the cerebellum is more evolutionarily conserved and primarily associated with basic brain functions. In contrast, the neocortex, acquired later in evolutionary time, is responsible for more advanced brain functions. Consequently, neurons such as TEGLU in the neocortex may be more vulnerable or dispensable compare to neurons in cerebellum, which serve primitive functions in aged mice. Nonetheless, the true biological significance of the dynamics between CBGRC and TEGLU necessitates further investigations.

In summary, we have shown that ChAIR effectively captures the dynamic changes in open chromatin loci, interactions, and RNA expression across cell-cycle phases in cultured cell lines and throughout various life stages in primary mouse brain cells. Our analyses indicate that the balance between short and ultra-long megacontacts in an individual cell’s genome correlates closely with its genomic properties and cell identity during its lifetime—including phases of maturation and aging. Additionally, the length of chromatin contacts correlates with genome folding density and nuclear volume in neuronal and non-neuronal cells, suggesting a potential nuclear mechano-genomic interplays that shape chromatin folding and accommodate transcriptional activity within the 3D nuclear space. With further development and refinement, the ChAIR strategy could be expanded to simultaneously detect both protein and RNA components that are crucial for maintaining and remodeling chromatin structures, epigenomic states, and transcription activities in a wide range of cells within complex biological systems.

---

**Fig. S1.**
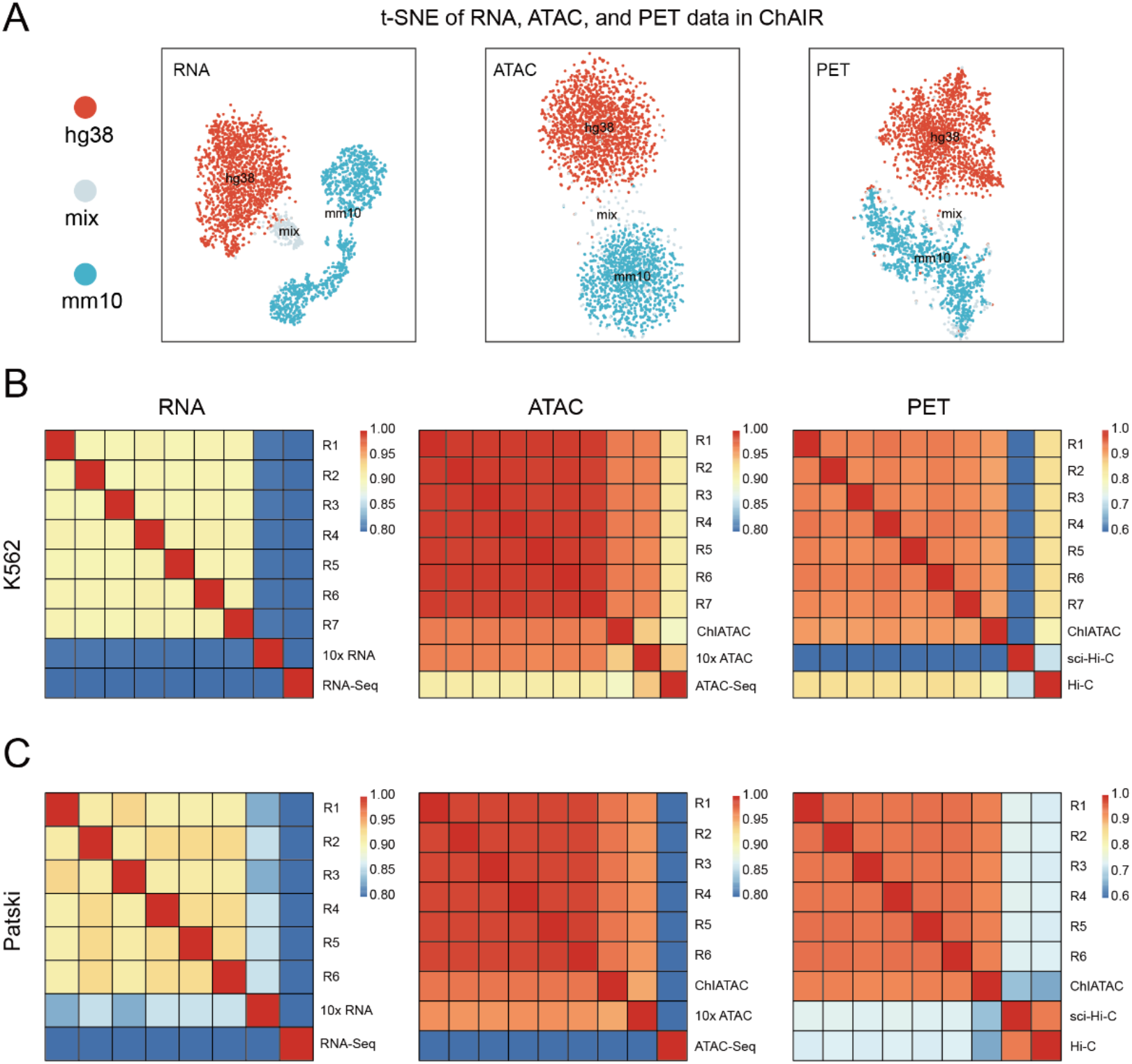
Quality assessment of ChAIR data from cell lines. (**A**) t-SNE plots of ChAIR-RNA, -ATAC, and -PET data from the barnyard experiment of human and mouse cell lines. (**B**-**C**) Correlation between technical replicates of ChAIR-RNA data in comparison with scRNA-seq from 10x Multiome (RNA+ATAC) and bulk RNA-seq data (left); ChAIR-ATAC data in comparison with ChIATAC, and scATAC-seq from 10x Multiome, and bulk ATAC-seq data (middle); ChAIR-PET data in comparison with ChIATAC, sci-Hi-C and bulk Hi-C data (right) in K562 (**B**) and Patski cells (**C**).

**Fig. S2.**
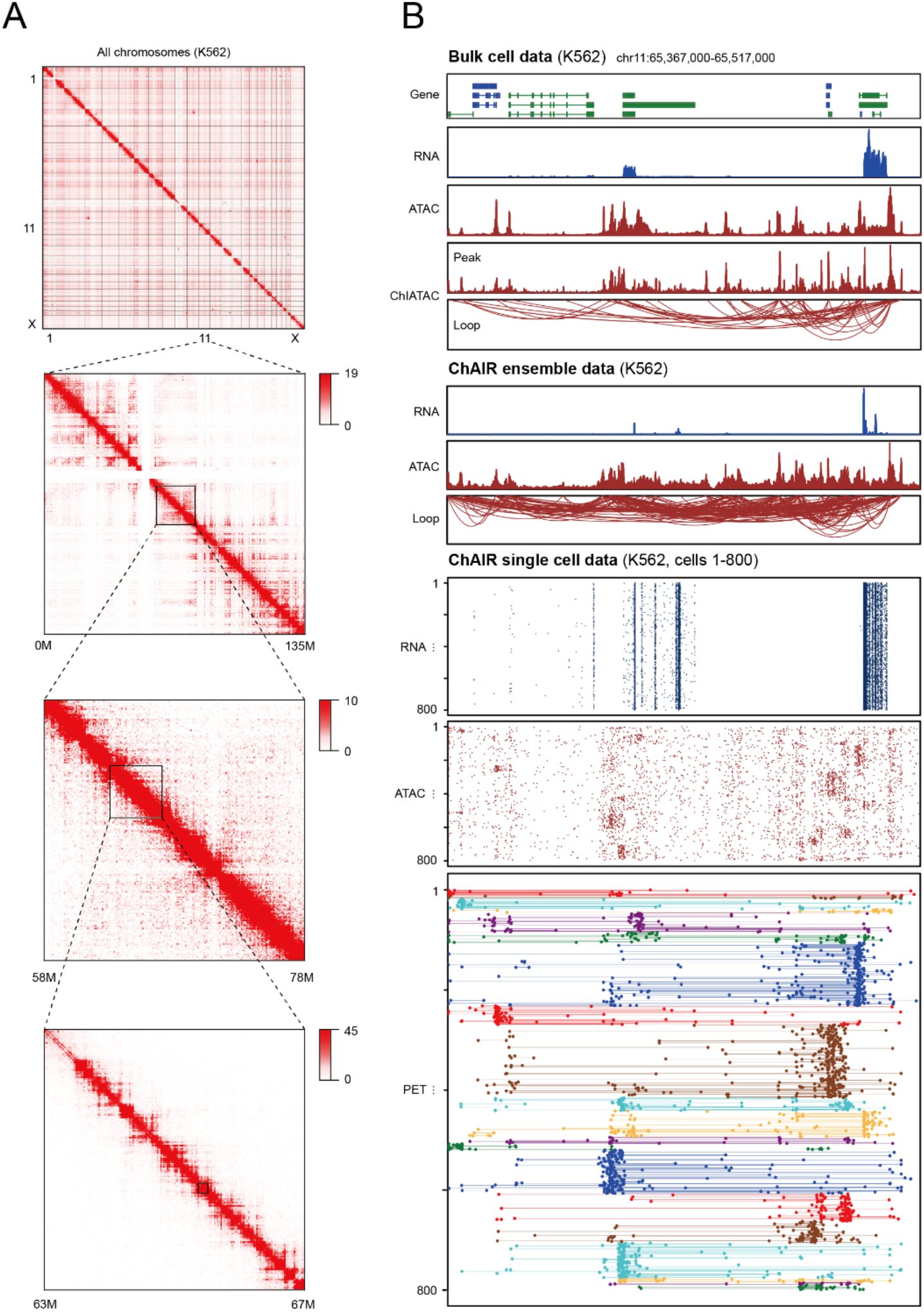
Additional example of ChAIR data visualization in K562 cells. (A) 2D contact heatmaps of ChAIR-PET data at varying resolutions. (B) A zoomed-in (150 kb in chr11) browser view of ChIATAC (loops and peaks), bulk cell ATAC-seq, and RNA-seq data (top) in comparison with ChAIR ensemble data (middle) and single-cell tracks of ChAIR-RNA, -ATAC, and -PET data (bottom).

**Fig. S3.**
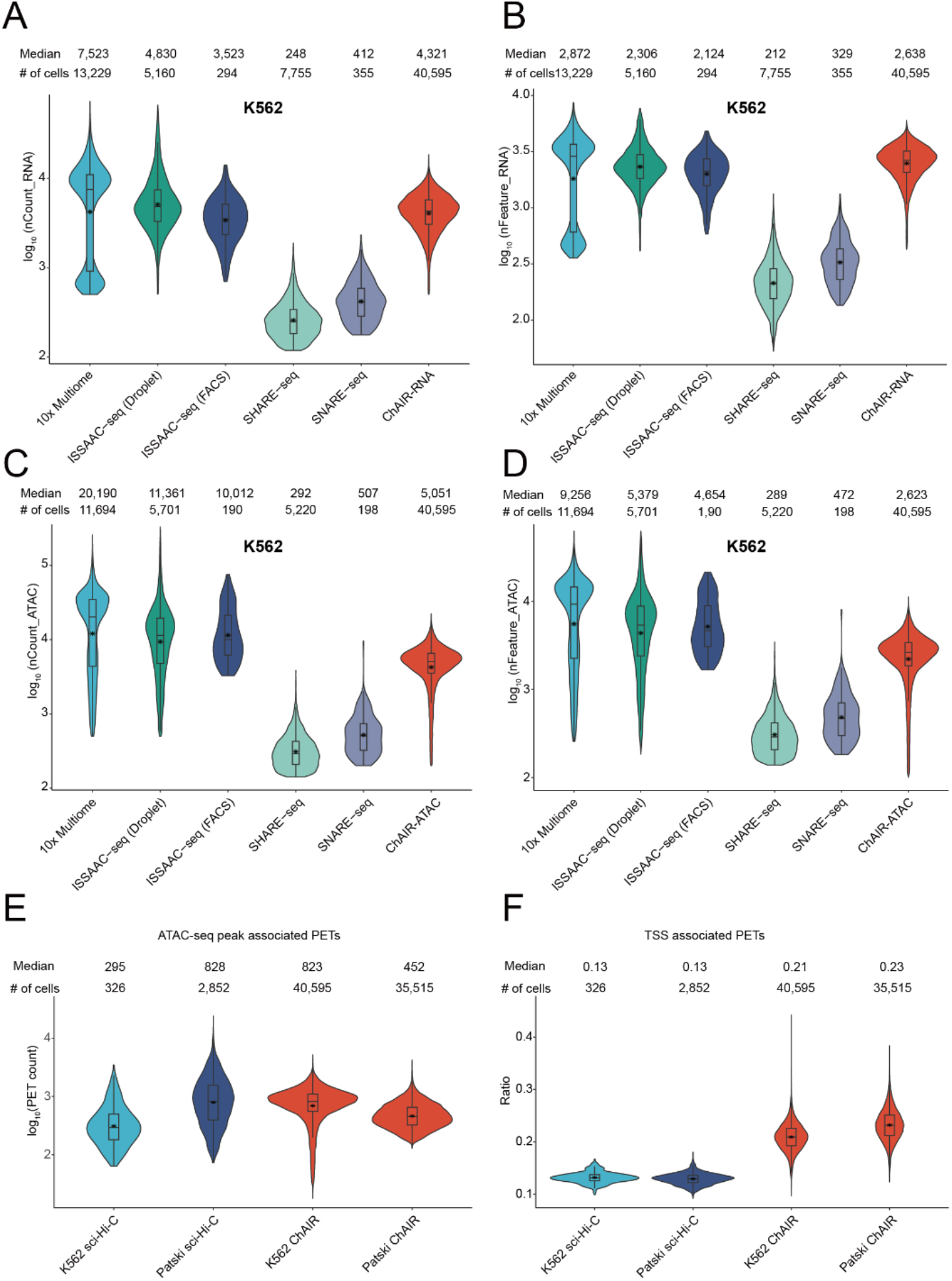
Benchmark ChAIR data derived from K562 and Patski cells with related methods. (A-D) Comparative analysis of multi-omic methods based on the following per-cell metrics: (A) number of RNA unique molecular identifiers (UMIs), (B) count of identified genes, (C) number of captured chromatin fragments, and (D) quantity of ATAC peaks. (E-F) Comparison between ChAIR and sci-Hi-C data in detecting chromatin contacts associated with ATAC peaks (E) and TSS sites (F). Medians and cell numbers in each category of data were provided.

**Fig. S4.**
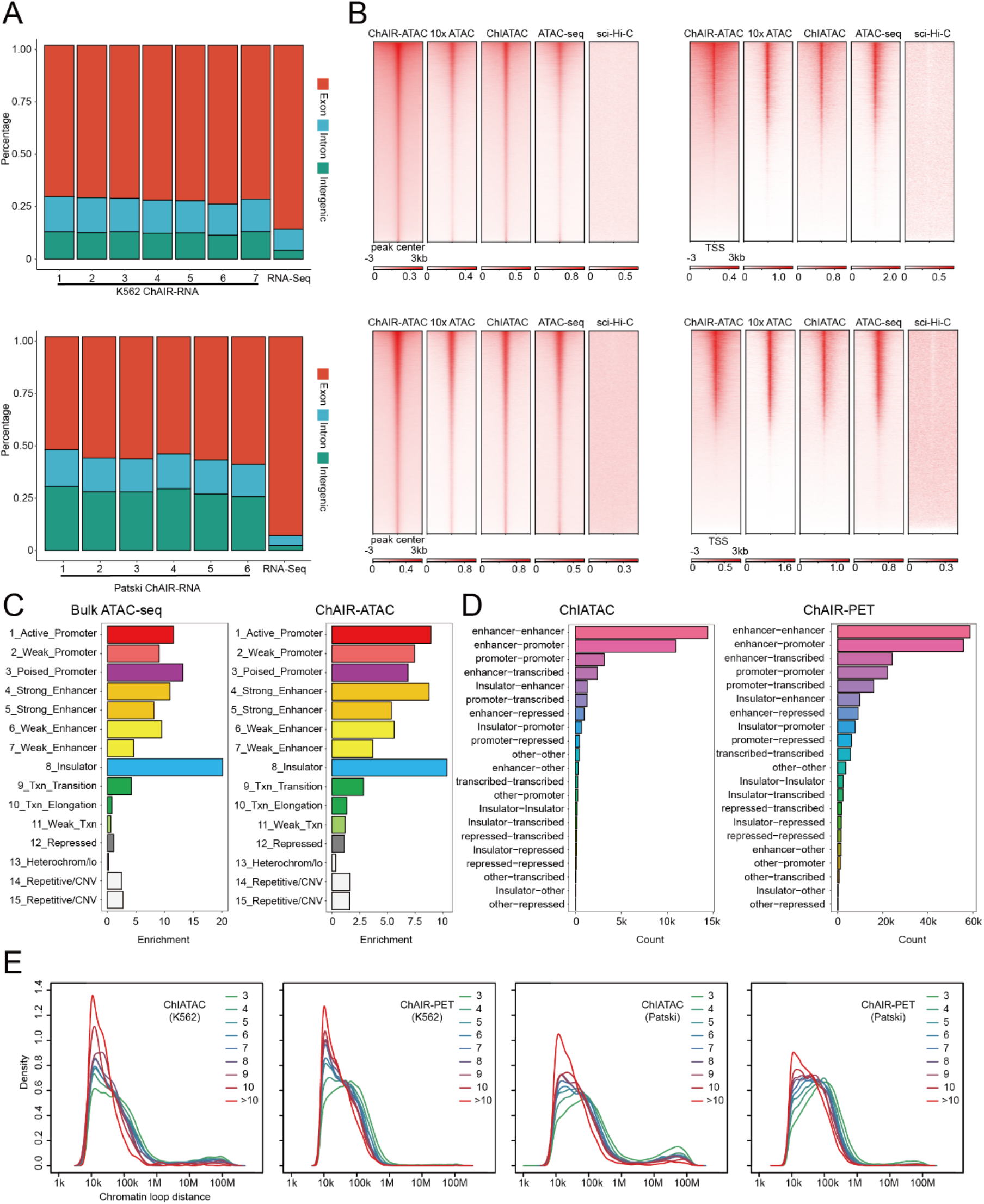
Characterization of ChAIR data. (**A**) Percentage of ChAIR-RNA reads that were mapped to exon, intron, and intergenic regions in K562 (top) and Patski cells (bottom) in reference with bulk RNA-seq data. (**B**) DNA mapping signals in ChAIR-ATAC peak loci (left) and at TSS sites (right) in comparison with 10x Multiome ATAC, ChIATAC, bulk ATAC-seq, and sci-Hi-C data in K562 (top) and Patski (bottom) cells. (**C**) Categories of ChromHMM defined chromatin states for open chromatin loci identified by bulk ATAC-seq data (n=176,891) (left) and ChAIR-ATAC data (n=201,540) (right) in K562. (**D**) Categories of ChromHMM defined chromatin states for chromatin loops in ChIATAC (n=37,579) (left) and ChAIR-PET data (n=232,988) (right). (**E**) Chromatin loop distance distribution of chromatin loops with different interaction frequency in ChIATAC and ChAIR-PET data in K562 and Patski cells.

**Fig. S5.**
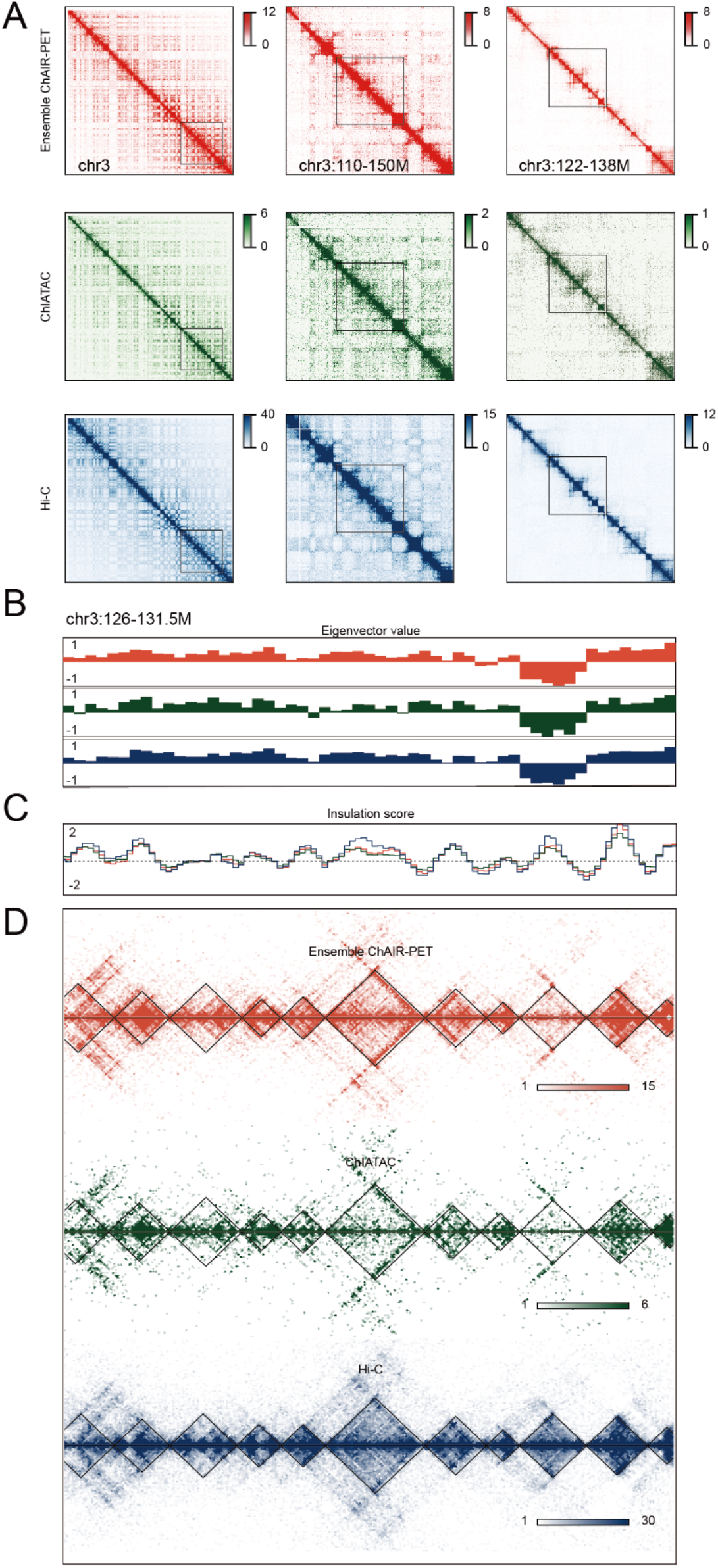
Comparison of chromatin contact profiles between ensemble ChAIR-PET, ChIATAC, and Hi-C data. (**A**-**D**) 2D contact heatmaps of chr3 at varying resolutions (**A**), Eigenvector values (**B**) and insulation scores (**C**), and TAD structures (**D**) identified in ensemble ChAIR-PET, ChIATAC and Hi-C data from Patski cells.

**Fig. S6.**
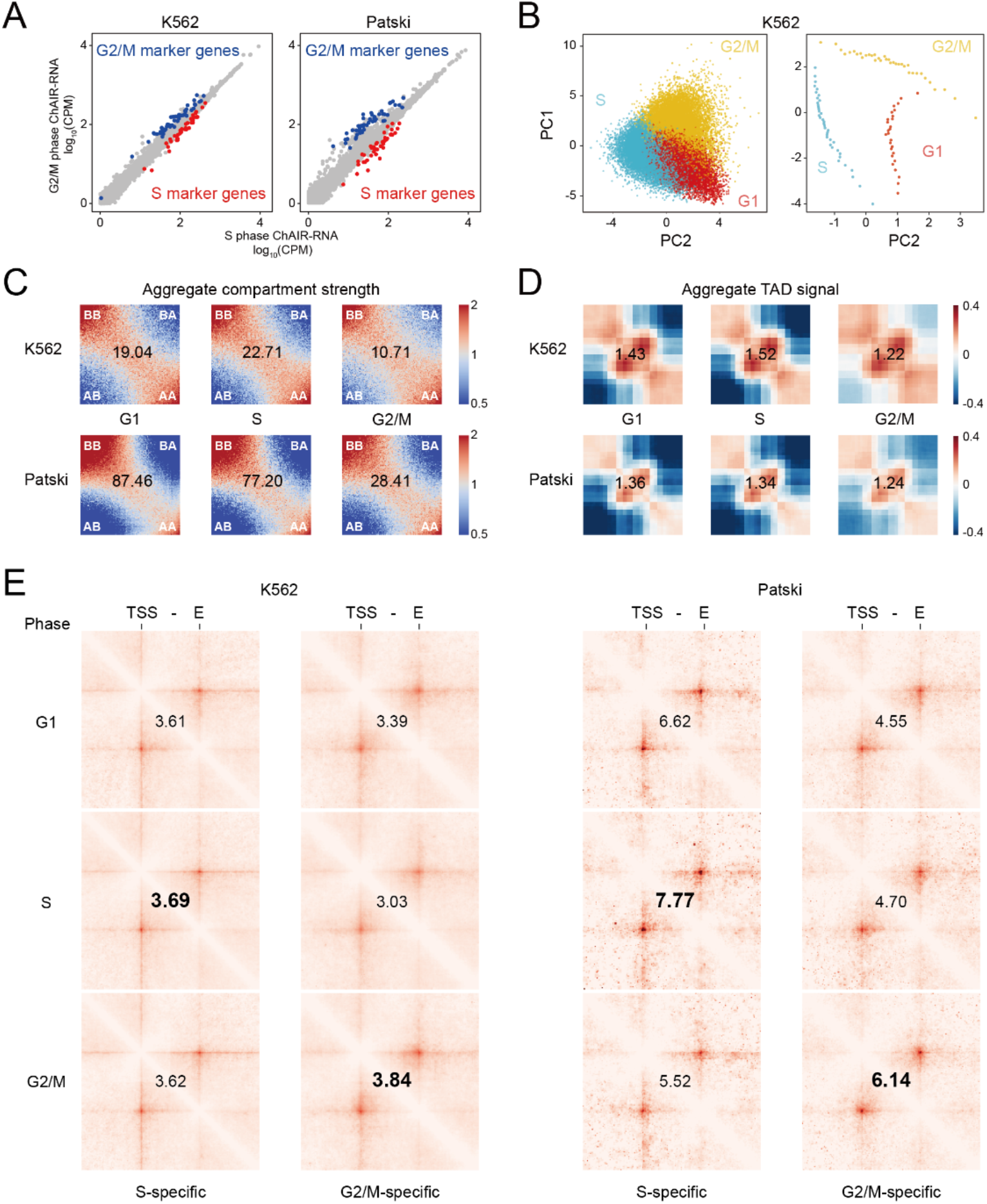
Cell-cycle specific profiles associated with cells phased by gene expression. (**A**) Scatter plots of gene expression of marker genes specific to S and G2/M phases ChAIR-RNA data. (**B**) PCA plots showing the distribution of individual K562 cells based on their cell-cycle marker genes (left) and further grouped into metacells based on cell-cycle pseudotime (right). (**C**) Aggregate compartment analysis of ChAIR-PET data in K562 (top) and Patski (bottom). (**D**) Aggregate TAD analysis of ChAIR-PET data in K562 (top) and Patski (bottom). (**E**) Aggregate plots of chromatin contacts between promoters (TSS) and putative distal enhancer elements (E) for phase-specific marker genes. The intensity values of phase-specific TSS-E interactions were given for comparison.

**Fig. S7.**
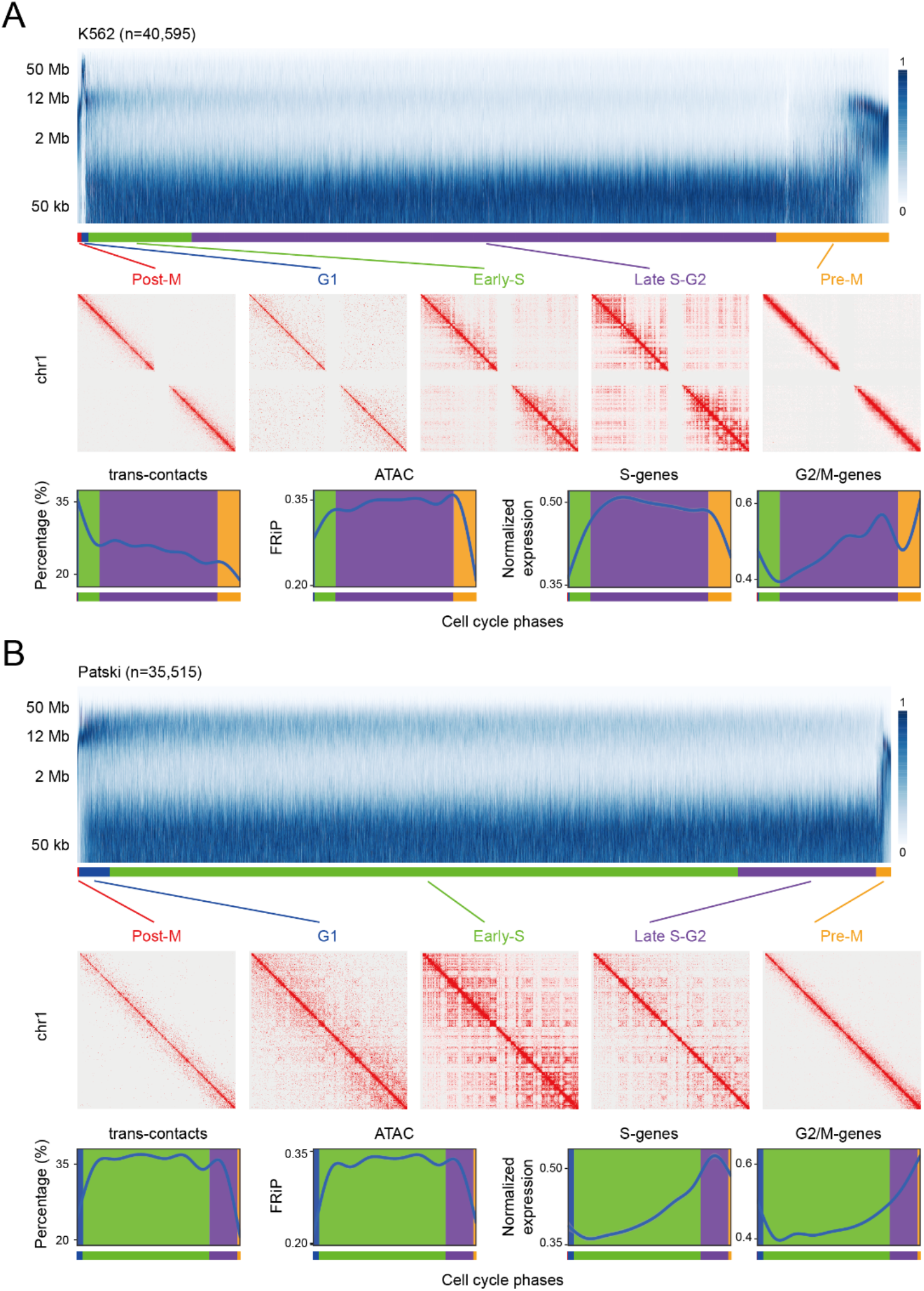
Spectrum of chromatin contact distances in correlation with cell-cycle specific profiles. (**A**-**B**) Chromatin contact distance distribution of ChAIR-PET data from K562 (**A**) and Patski (**B**) cells. Individual cells were sorted based on the extent of chromatin interactions within two specific ranges: 20kb-1Mb and 2-12Mb (top, see methods). Cells were defined as in post-Mitosis, G1, Early S, Later S-G2, and pre-Mitosis phases with characteristic chromatin folding patterns (middle). The profiles of the percentage of inter-chromosomal (trans) chromatin contacts in ChAIR-PET data, fraction of reads in peak (FRiP) in ChAIR-ATAC data, and gene expression of cell cycle marker genes in ChAIR-RNA data in each individual cells in corresponding to cell cycle phases were also shown (bottom).

**Fig. S8.**
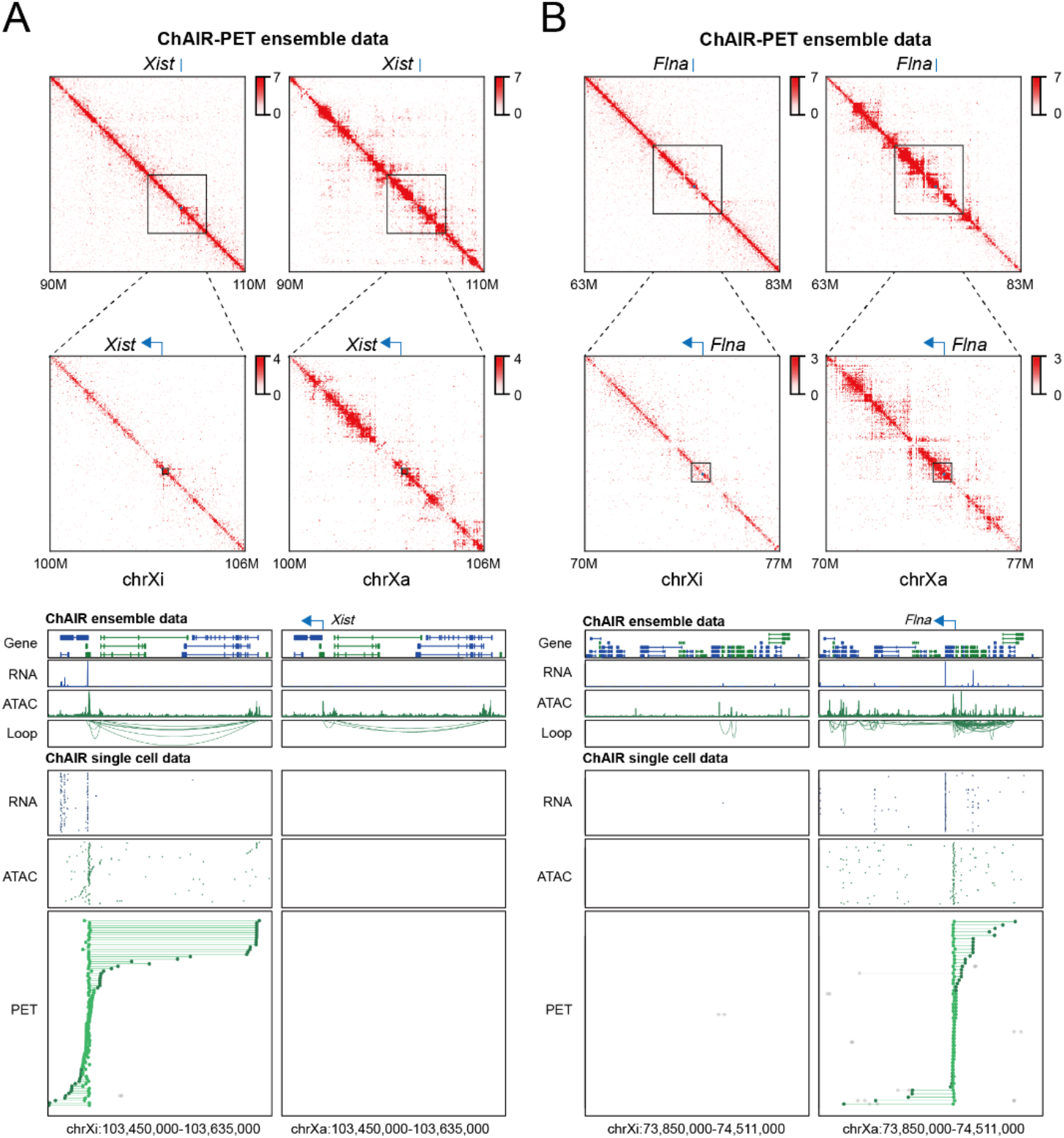
Allelic-specific chromatin interaction, accessibility, and gene expression in Patski cells. Example views of data visualization at the loci of *Xist* (**A**) and *Flna* (**B**). 2D contact heatmaps at varying resolutions surrounding active chrX (chrXa) and inactive chrX (chrXi) in ChAIR-PET data (top), and the browser views of ensemble and single-cell tracks of ChAIR-RNA, -ATAC, and -PET data (bottom) were shown.

**Fig. S9.**
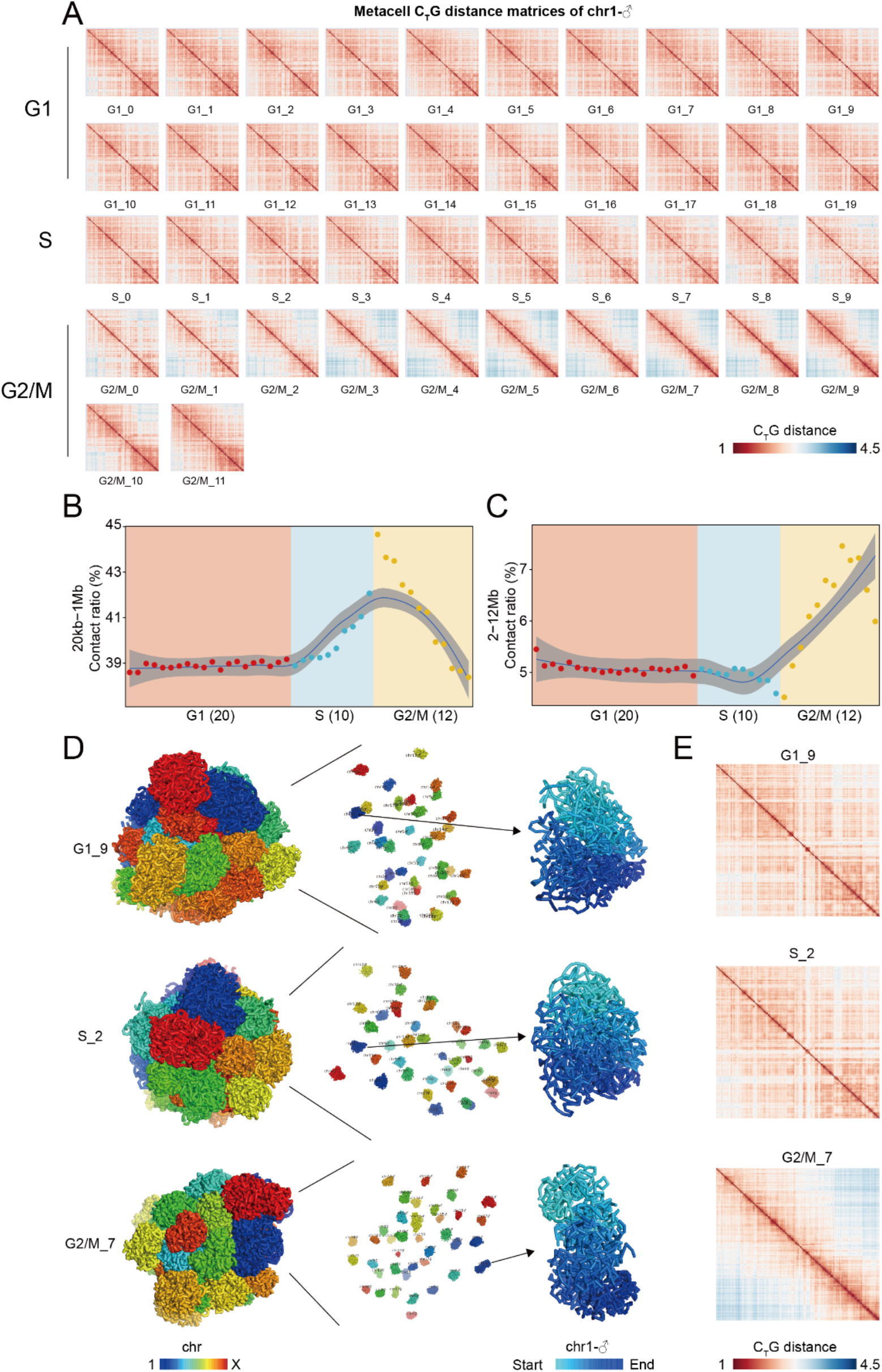
Haplotype-resolved chromatin features during cell cycle. (**A**) C_T_G distance matrices for the paternal copy of chr1 across 42 metacells in 3 cell cycle phases. (**B**-**C**) The percentage of chromatin contacts in the range of 20kb-1Mb (**B**) and 2-12Mb (**C**) in cell cycle phases. (**D**) Representative 3D models of the nuclear architectures from 3 specific metacells with an expanded view that includes haplotype-resolved chromosomes and an enlarged view of chr1-♂. (**E**) The corresponding C_T_G distance matrices of specific metacells as shown in (**D**).

**Fig. S10.**
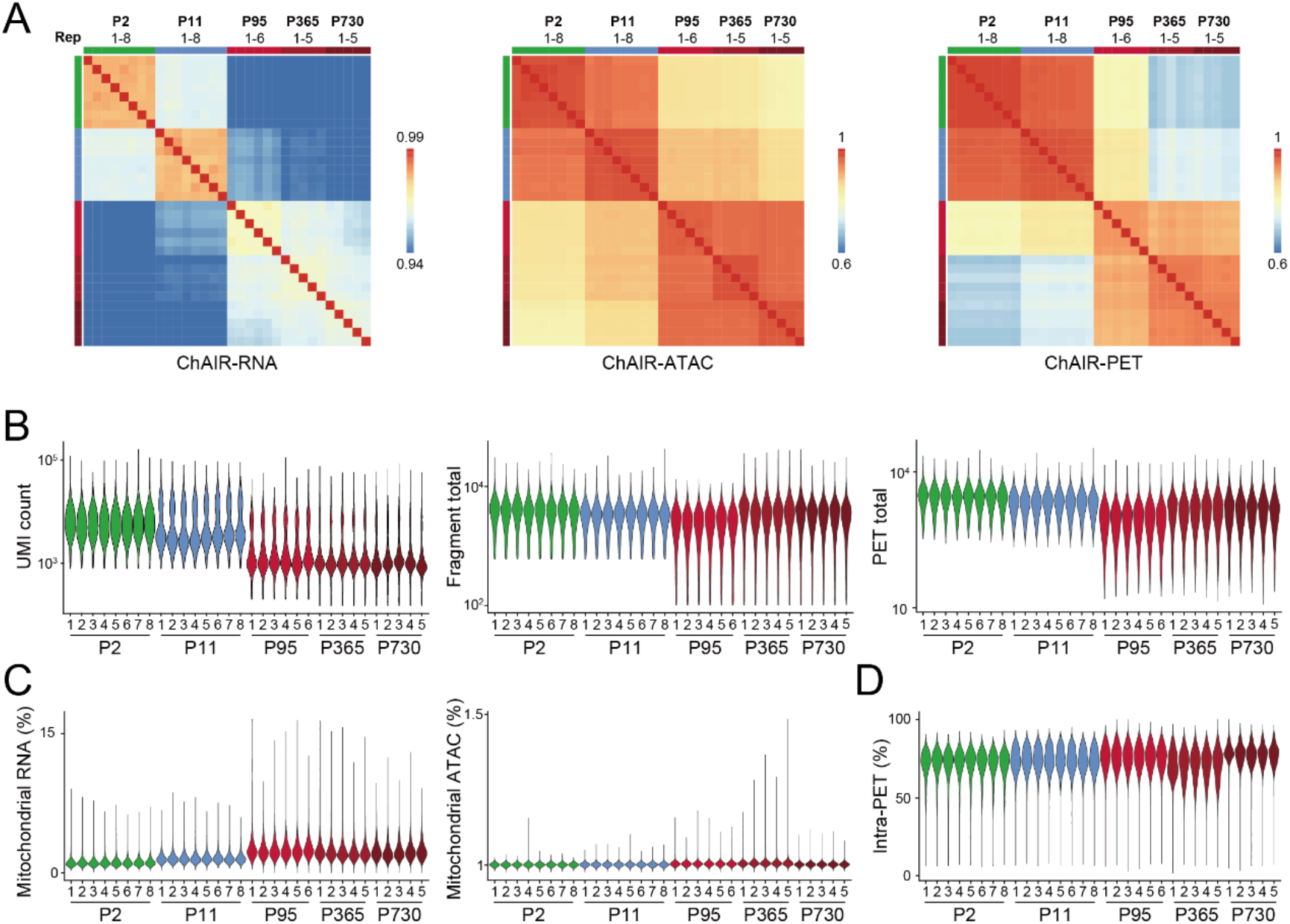
Reproducibility assessment of ChAIR data from mouse brains. (**A**)Reads correlation between technical replicates from ChAIR-RNA, -ATAC, and -PET data. (**B**) Violin plots showing the counts of UMIs, fragments, and PETs captured in ChAIR data. (**C**) Percentage of mitochondrial RNA and mitochondrial DNA reads in ChAIR-RNA and -ATAC data. (**D**) Percentage of intra-chromosomal PET (Intra-PET) in ChAIR-PET data.

**Fig. S11.**
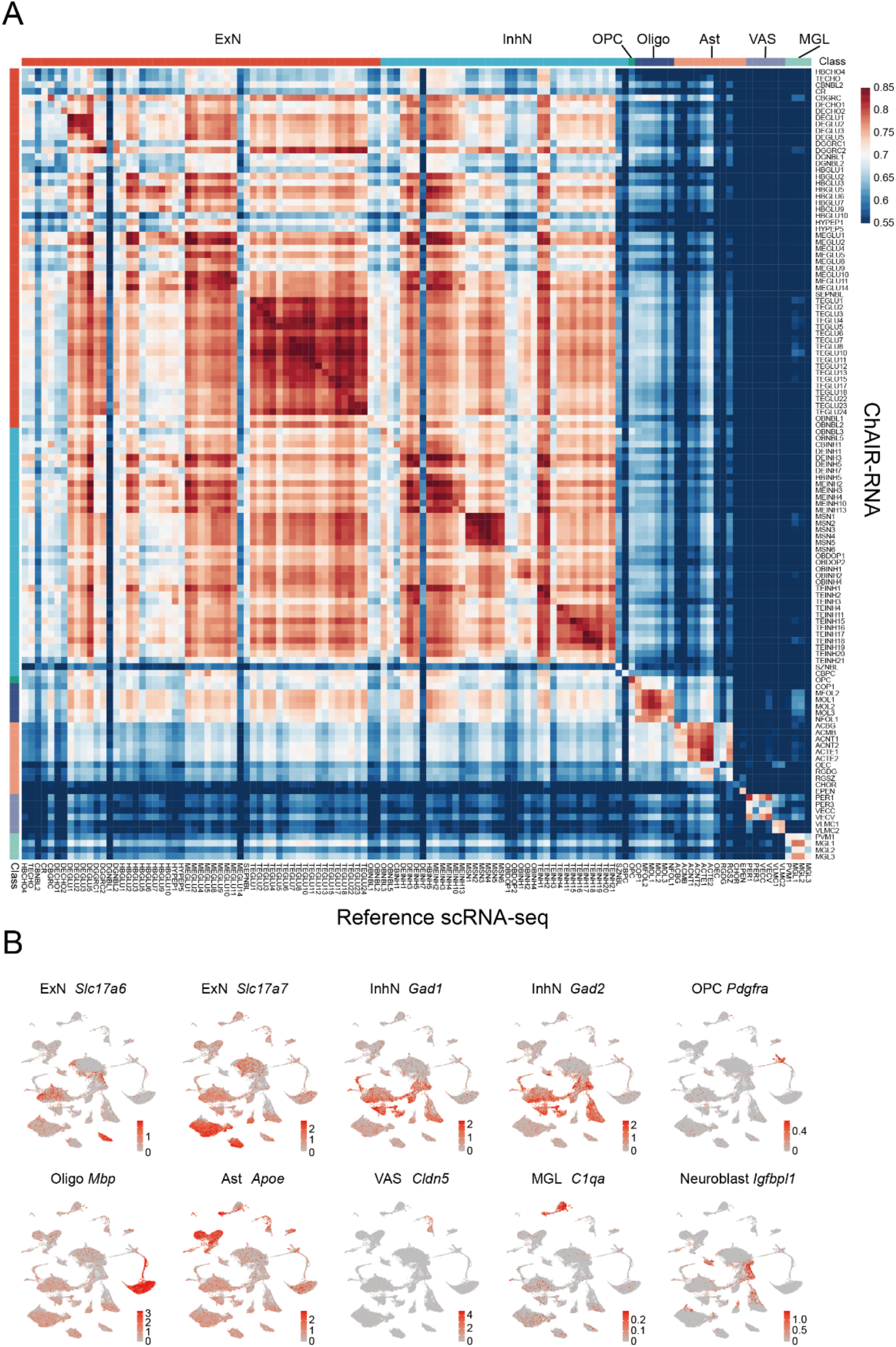
Quality assessment of ChAIR-RNA data from mouse brain cells. (**A**) Correlation between ChAIR-RNA and a reference scRNA-seq data. (**B**) UMAP plots showing the expression signals of canonical marker genes in ChAIR-RNA data.

**Fig. S12.**
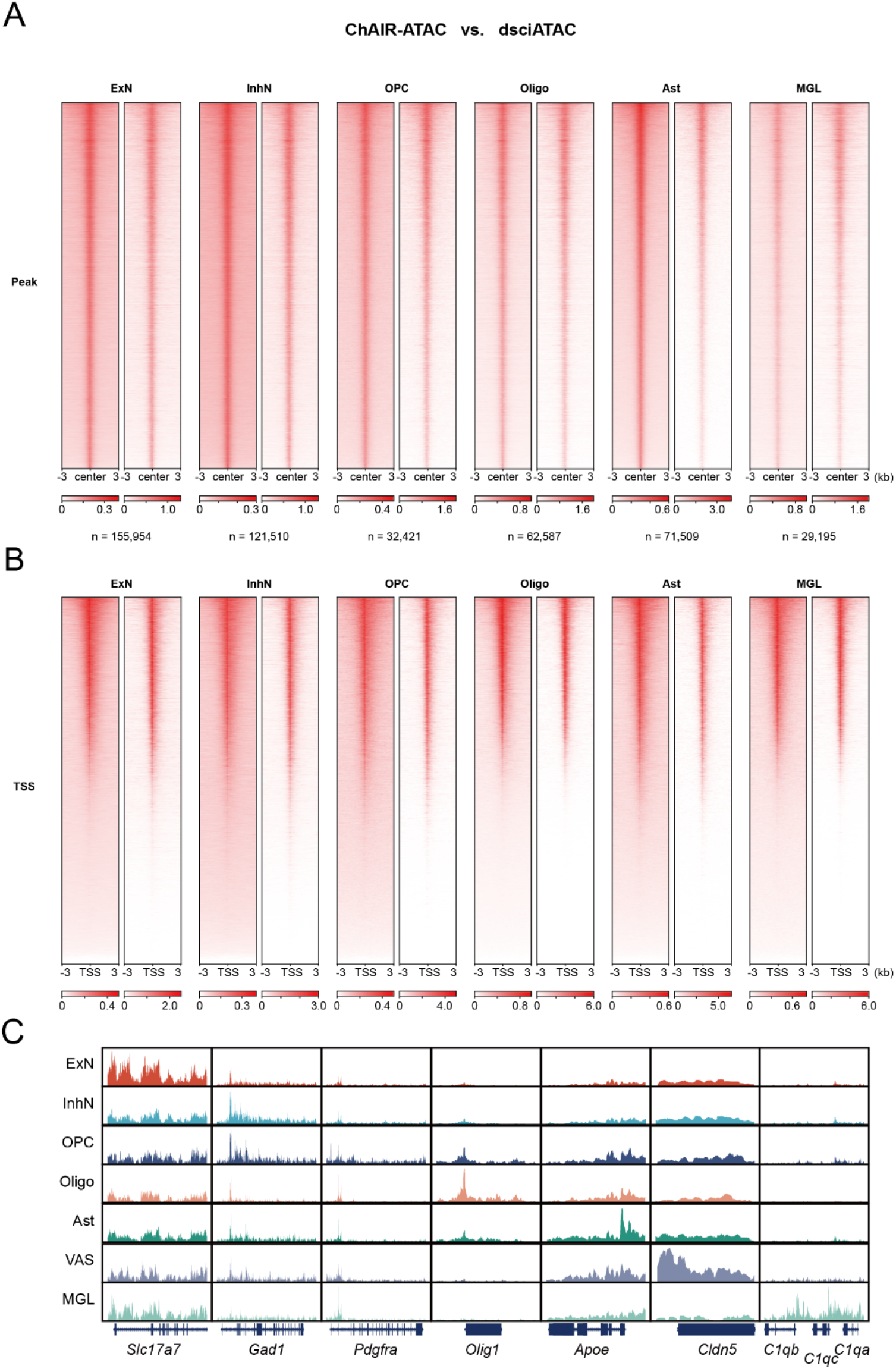
Quality assessment of ChAIR-ATAC data from mouse brain cells. (**A**-**B**) Side-by-side comparison of signal enrichment at ATAC peak loci (**A**) and TSS loci (**B**) between ChAIR-ATAC and dsciATAC data across major brain cellular classes. Numbers of ATAC peaks (n) for each class were given. (**C**) Browser view examples of ensemble ChAIR-ATAC signals at marker gene loci across different cellular classes.

**Fig. S13.**
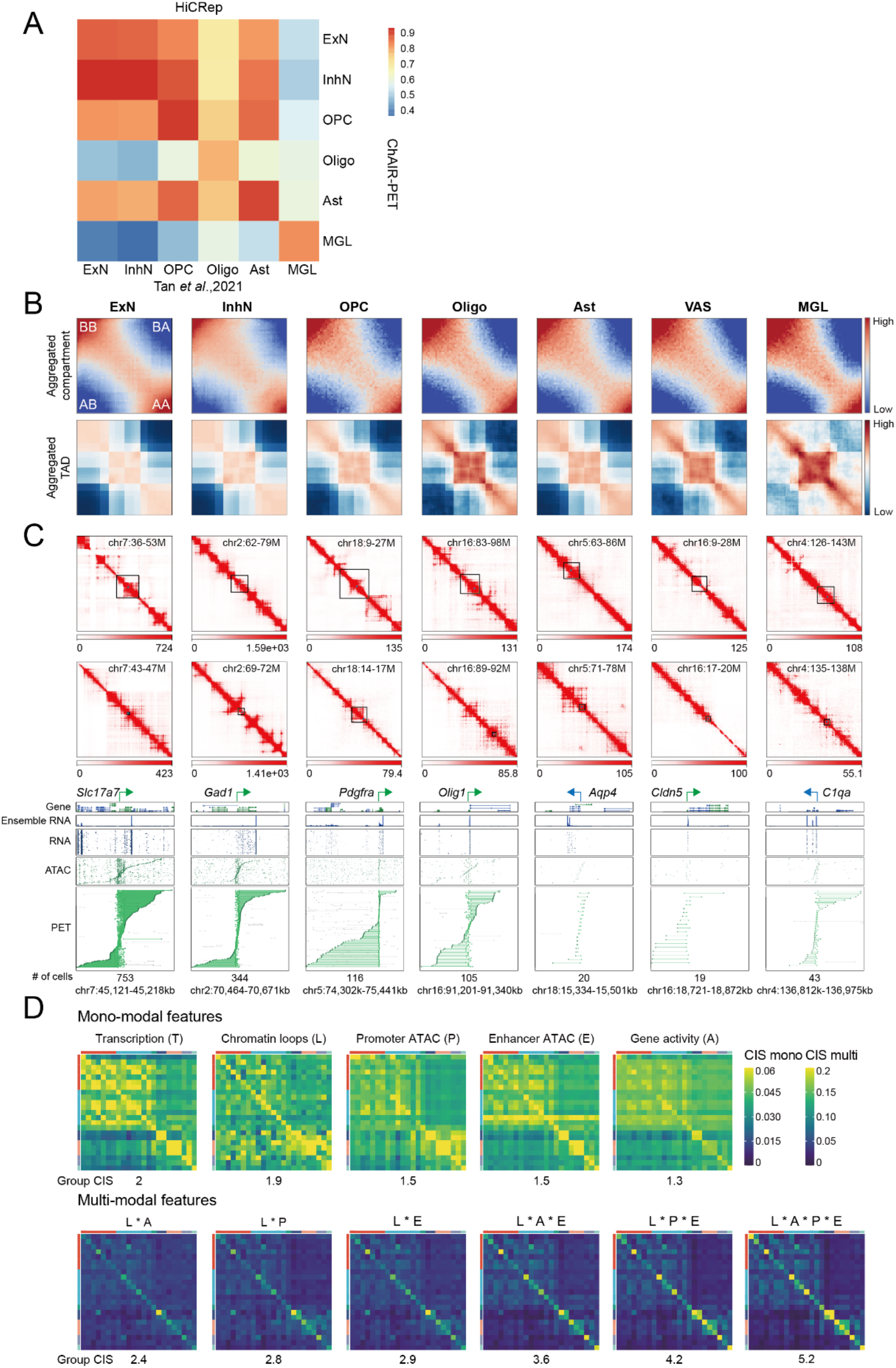
Quality assessment and characterization of ChAIR-PET data from mouse brain cells. (**A**) HiCRep analysis comparing cell-specific chromatin contact profiles between dataset from Tan *et al.*, 2021, and ChAIR-PET data. (**B**) Aggregate chromatin compartment (top) and TAD (bottom) analysis of ChAIR-PET data in 7 cellular classes. (**C**) Example views of 2D contact heatmaps at varying resolutions around specific marker genes (top). Corresponding browser views of single-cell tracks of ChAIR-RNA, -ATAC, and -PET data were also shown (bottom). (**D**) Cell identity score (CIS, see methods) matrices calculated using different mono-modal features associated with marker genes including transcription (T), ATAC signals for gene body (Gene activity, A), promoter (P), enhancer (E), and chromatin loop (L), and multi-modal 3D epigenomic features in various combination across 23 cell groups. The value of group CIS for each feature reflects its specificity in defining cell identity.

**Fig. S14.**
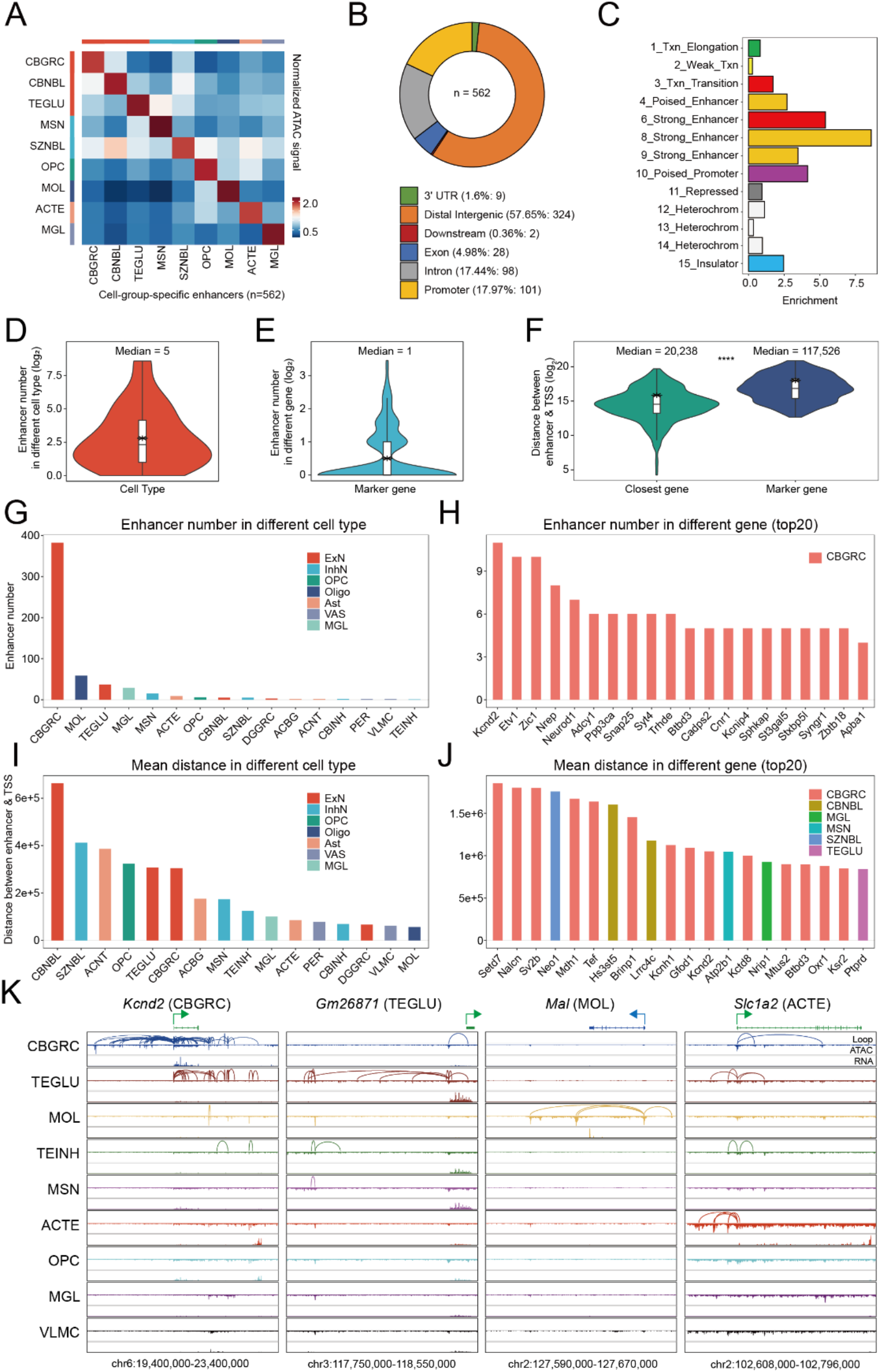
Characterization of cell-specific enhancers. (**A**) The matrix of ATAC signals at cell-specific enhancer loci (n=562) across 9 cell groups. (**B**) Genomic annotations of cell-specific enhancers. (**C**) Chromatin states of cell-specific enhancers annotated by ChromHMM. (**D**-**E**) Number of enhancers in different cell groups (**D**) and target marker genes (**E**). (**F**) The distance between enhancers and both nearby and target genes. (**G**-**H**) Numbers of enhancers identified in each cell group (**G**) and associated with each target gene (**H**). (**I**-**J**) Mean distance of enhancer to target gene’s TSS within different cell groups (**I**) and for individual genes (**J**). (**K**) Examples of cell-specific enhancers across 9 cell groups.

**Fig. S15.**
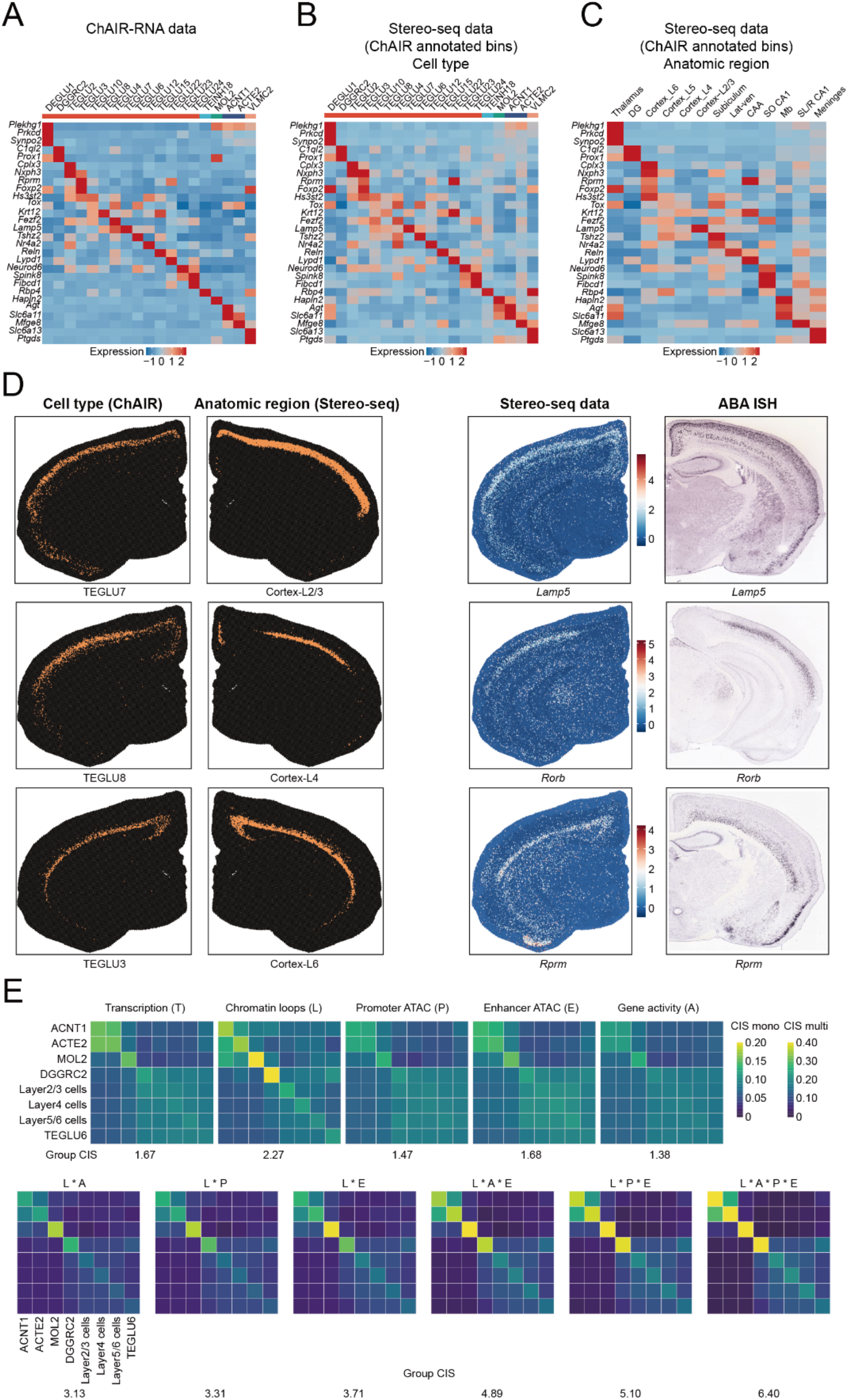
Validation of integrating ChAIR and Stereo-seq data. (**A**-**C**) Canonical marker gene expression profile in ChAIR-RNA (**A**) and Stereo-seq data (**B**-**C**). Stereo-seq data with cell type annotations (**B**) and anatomic region annotations (**C**) were shown. (**D**) Spatial visualization of cells from various cerebrum cortex layers and the gene expression of marker genes in Stereo-seq and Allen Brain Atlas in situ hybridization (ABA ISH) data. (**E**) CIS matrices calculated by different features across region-resolved cell types.

**Fig. S16.**
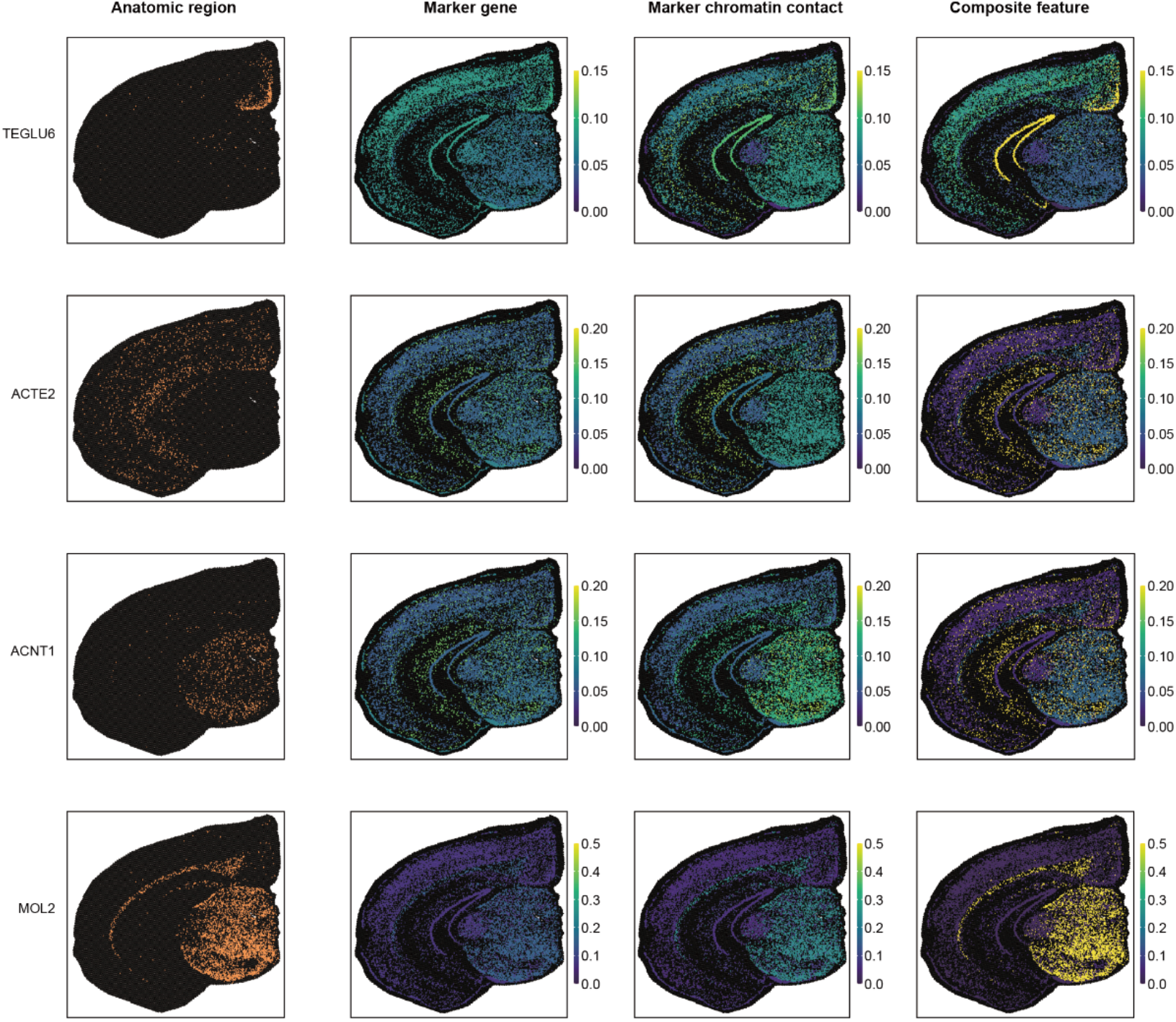
Additional examples of regional profiles of spatially-resolved ChAIR data.

**Fig. S17.**
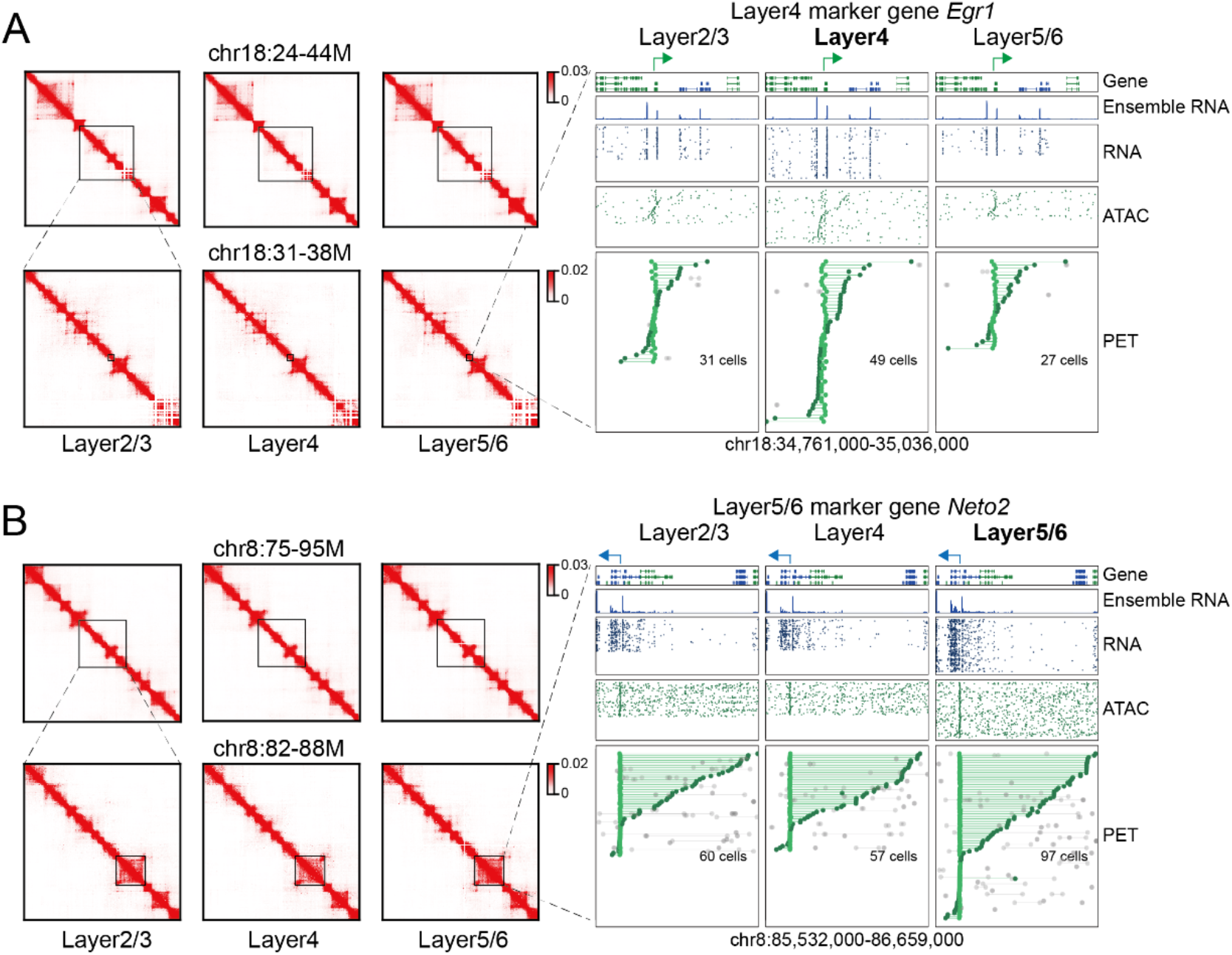
Additional examples of ChAIR data showing chromatin features at cortex layer-specific marker genes. (A) ChAIR data showing ensemble 2D contact heatmaps (left), and single-cell tracks of marker gene promoter associated chromatin contacts (PET), ATAC, and RNA at cortex layer4 marker gene *Egr1* locus (right). Similar analysis was done for cortex layer5 marker gene *Neto2* (B).

**Fig. S18.**
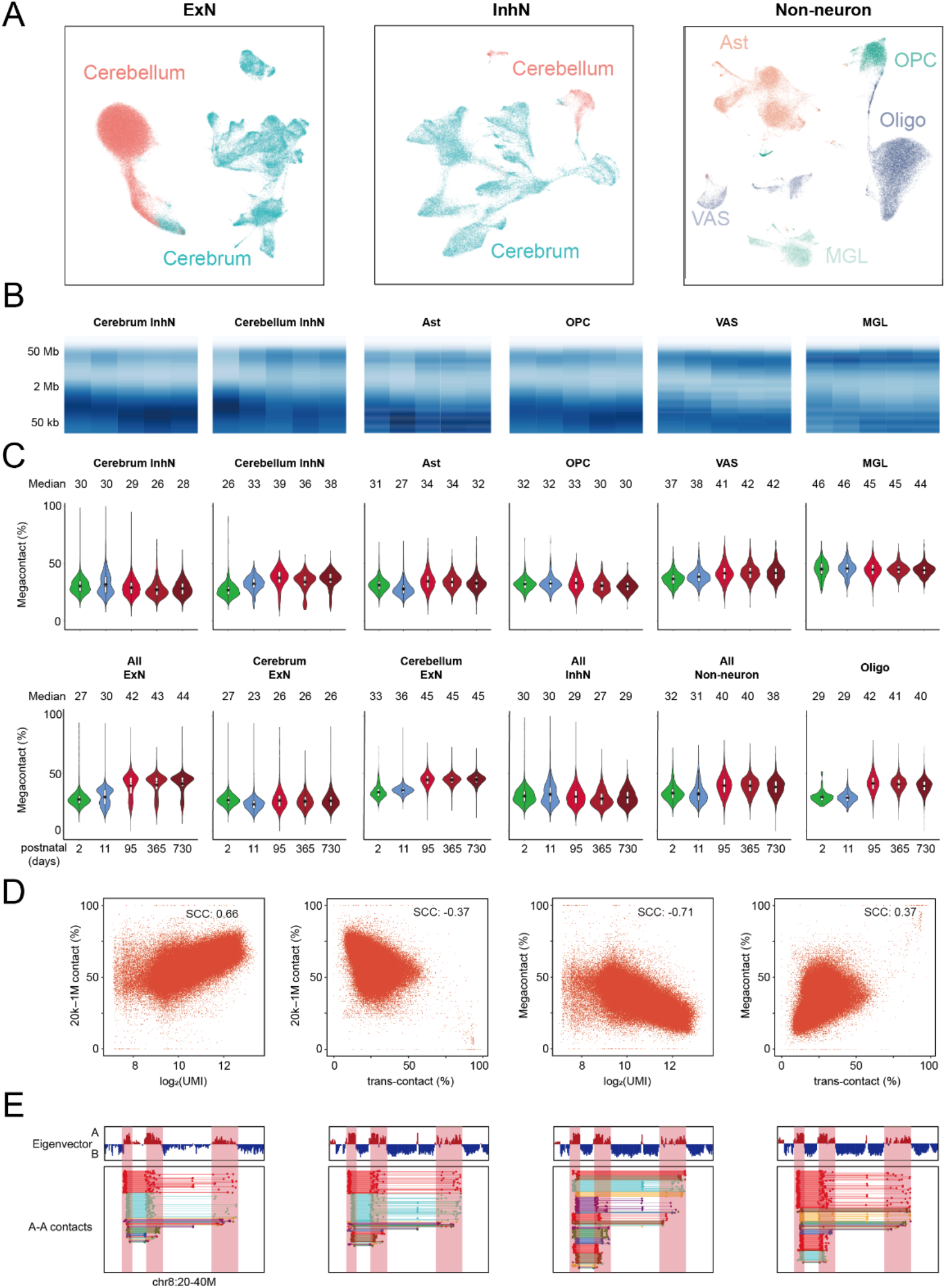
Dynamics of 3D epigenome of mouse brain cells during aging. (**A**) UMAP plots of re-embedded ChAIR-RNA data of ExNs, InhNs, and non-neuronal cells. (**B**) Chromatin contact distance spectrum in different cells across 5 age points. (**C**) The percentage of ultra-long megacontacts (>2Mb) in different cells. (**D**) The correlation between the extent of short-range contacts (20kb-1Mb) (left) and the ultra-long megacontacts (right) with global transcription activity and trans-contact. (**E**) Single-cell browser views of ChAIR-PET data showing chromatin contacts between nearby A compartments.

**Fig. S19.**
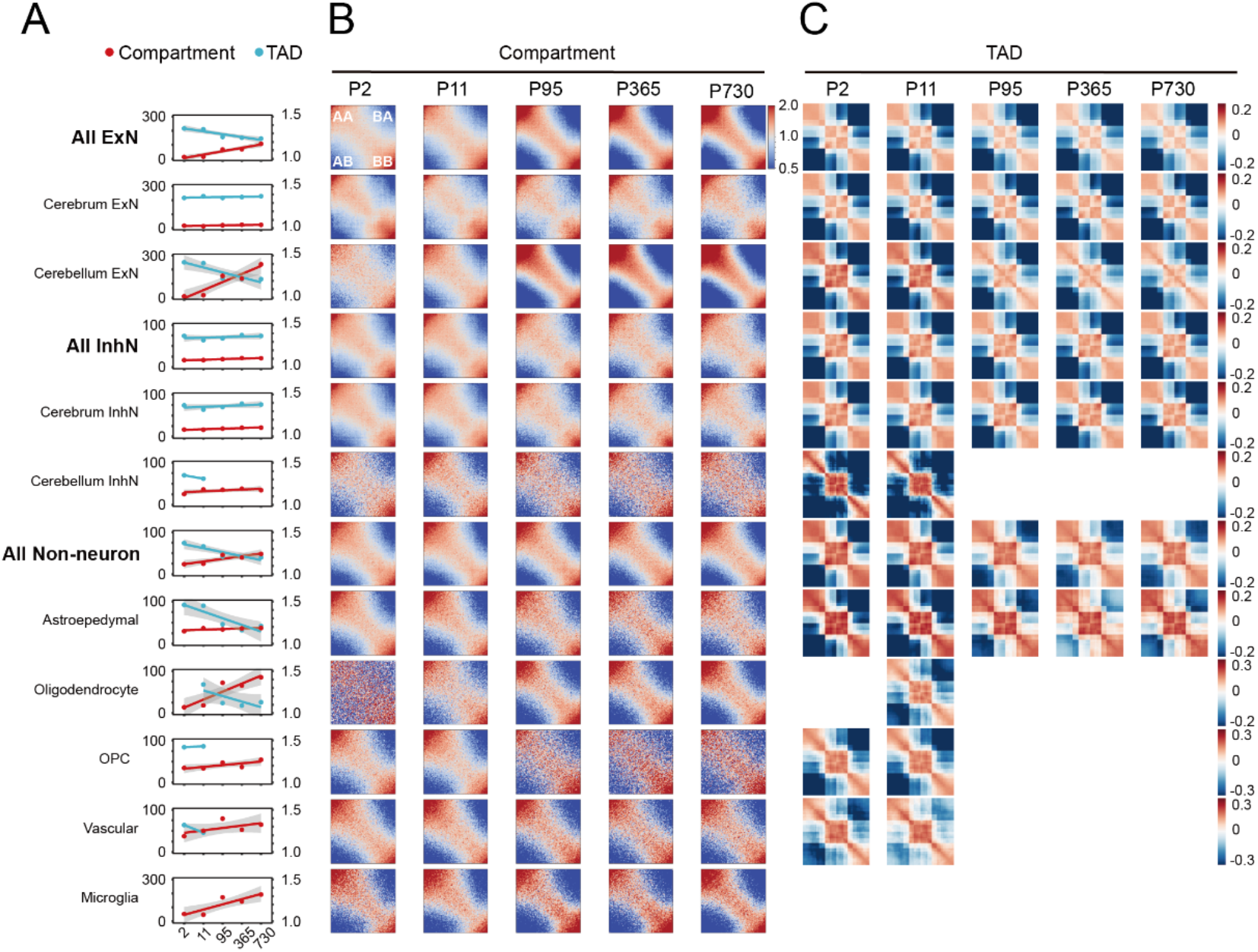
The dynamics of chromatin features in mouse brain cells during aging. (A-C) Summary of chromatin features (A) including compartment strength (B) and aggregated TAD signals (C) across various cell types during aging.

**Fig. S20.**
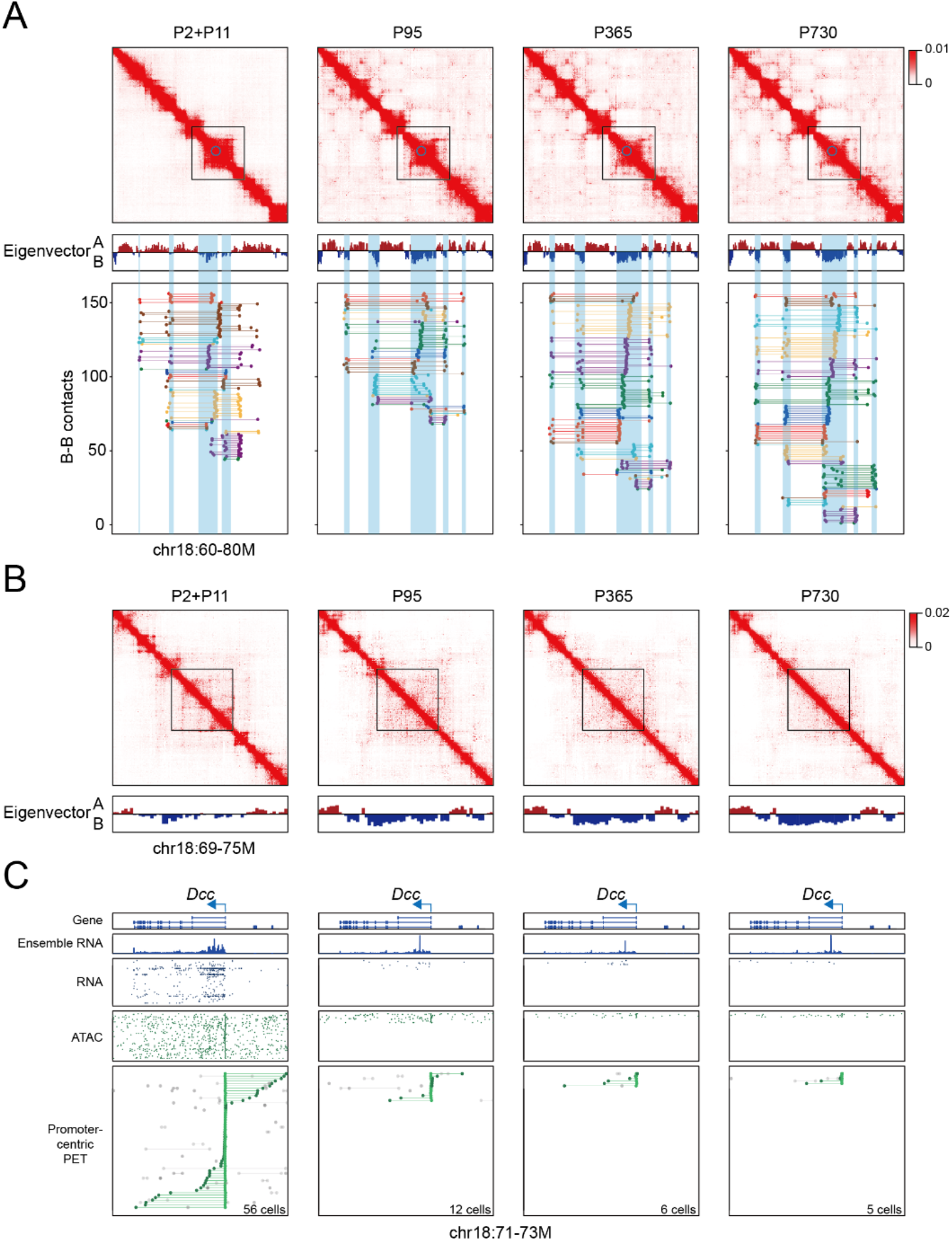
Additional example showing the concurrent changes of chromatin architecture and transcription during aging. (**A**) ChAIR-data showing ensemble 2D contact heatmaps, eigenvector values, contacts between B compartments in single cells. (**B**) Zoom-in views of boxed regions of 2D contact heatmaps from (**A**), showing the disappearance of TAD structures during aging. (**C**) Marker gene promoter-associated chromatin contacts at the *Dcc* locus in CBGRC.

**Fig. S21.**
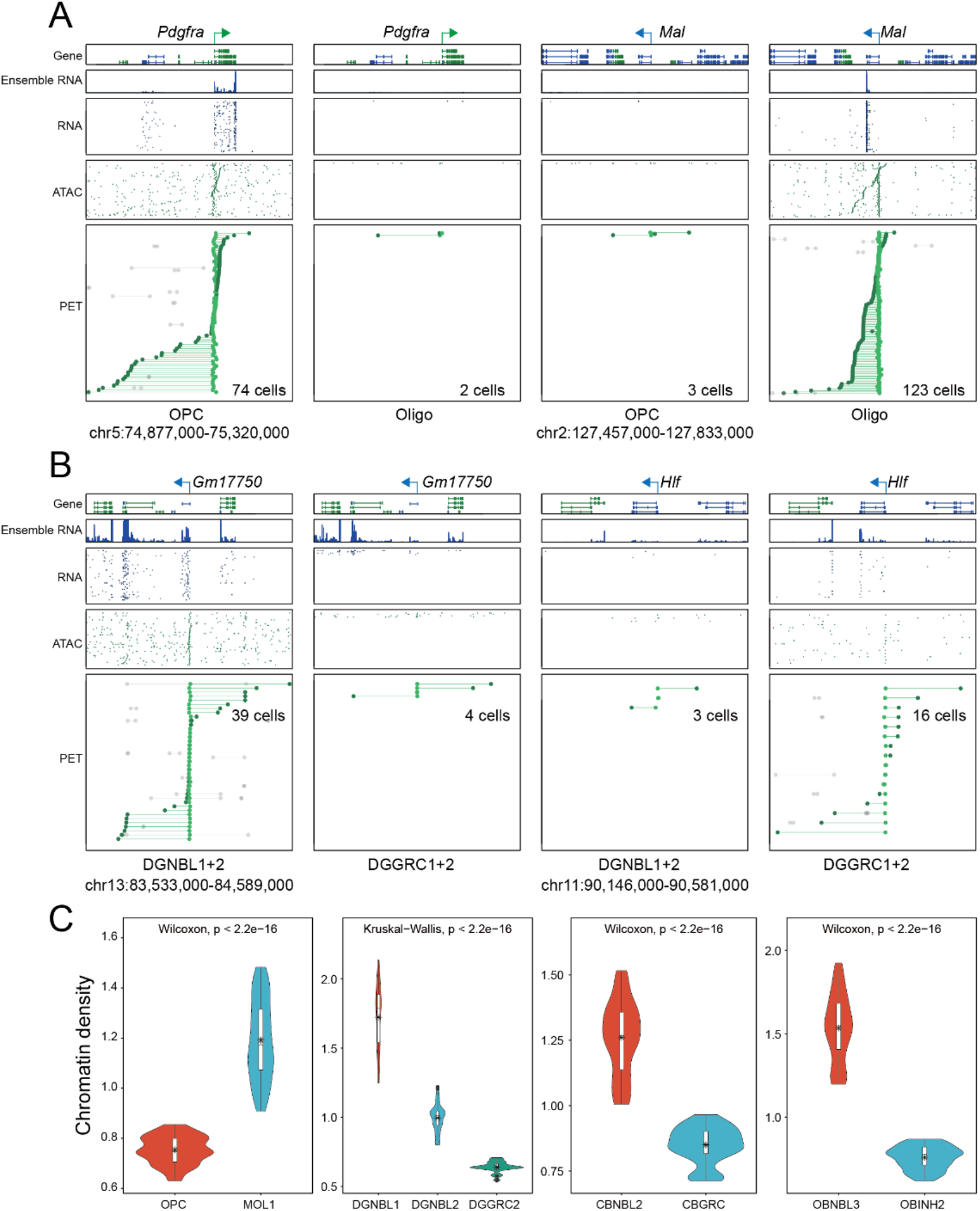
Dynamics of chromatin architecture and transcriptome during cell differentiation. (**A**-**B**) Single-cell tracks of ChAIR data showing promoter-associated chromatin contacts of marker genes in OPC and Oligos (**A**) and ExNs in DG (**B**). (**C**) The measurement of chromatin density estimated from the 3D models of nuclear architecture derived from metacells of non-neuronal cells (OPC and MOL1) and neurons in DG, CB, OB.

**Fig. S22.**
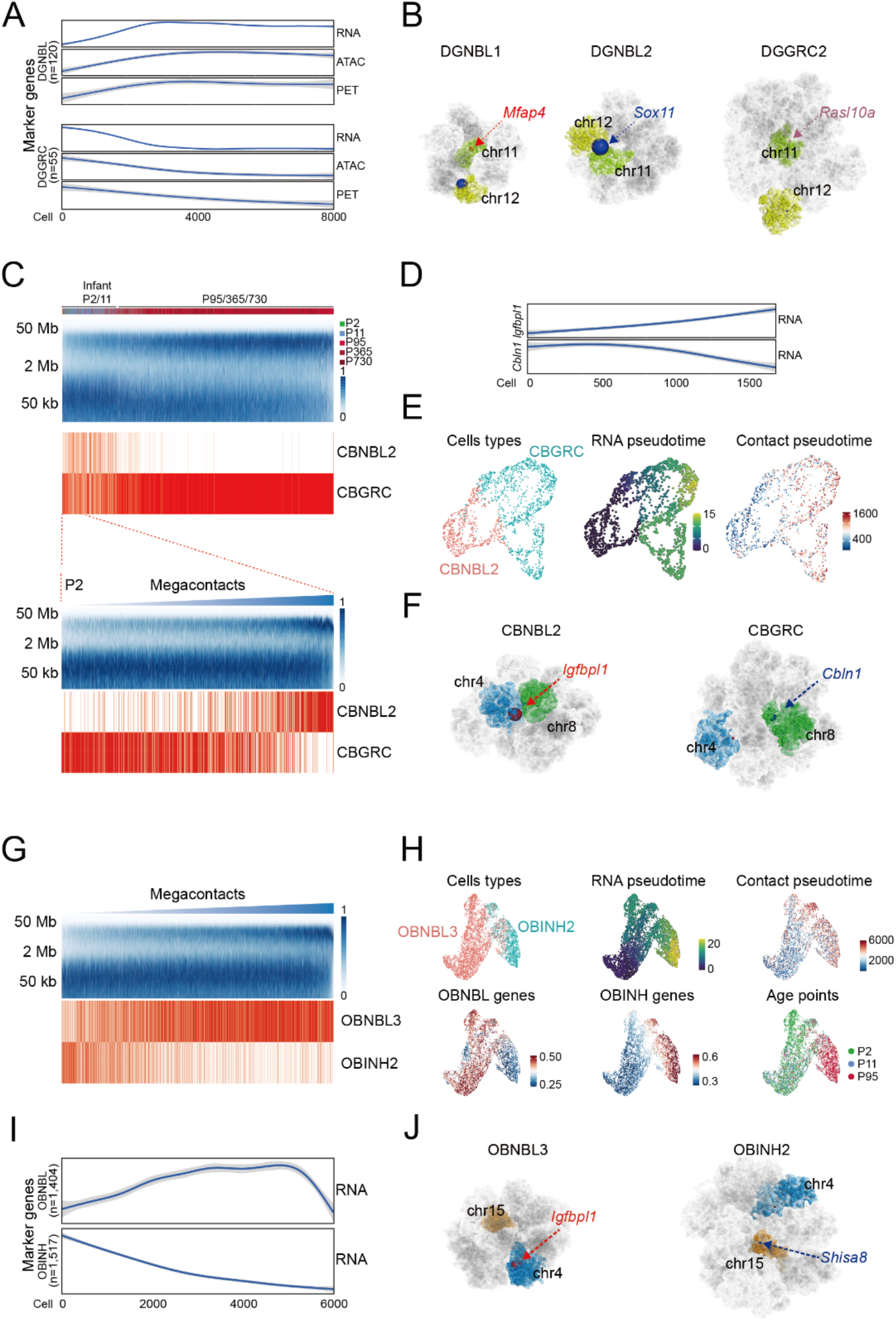
Examples of chromatin rewiring in neurons during differentiation. (**A**) Correlation of chromatin contact distances with gene expression (RNA), promoter’s chromatin accessibility (ATAC) and connectivity (PET) in individual cells. Cells were sorted by the extent of megacontacts, from lowest to highest. (**B**) The reconstructed 3D models of nuclear architectures from DGNBL1, DGNBL2, and DGGRC2 metacells. Nuclear positions of specific marker genes were indicated. The sizes of balls reflected the relative levels of gene expression. (**C**) Contact distance spectrum of cerebellar ExNs (CBNBL2 and CBGRC) sorted from the least extent of long-range chromatin contact (>2Mb) to the highest (top), with an additional focus on the P2 stage (bottom). (**D**) Chromatin contact pseudotime analysis of P2 ChAIR data. (**E**) UMAP plots of ChAIR data with RNA and chromatin contact pseudotime information in P2 ChAIR data. (**F**) 3D models of nuclear architectures reconstructed from CBNBL2 and CBGRC at P2 stage. Similar analyses were done in InhNs from OB (**G**-**J**).

**Fig. S23.**
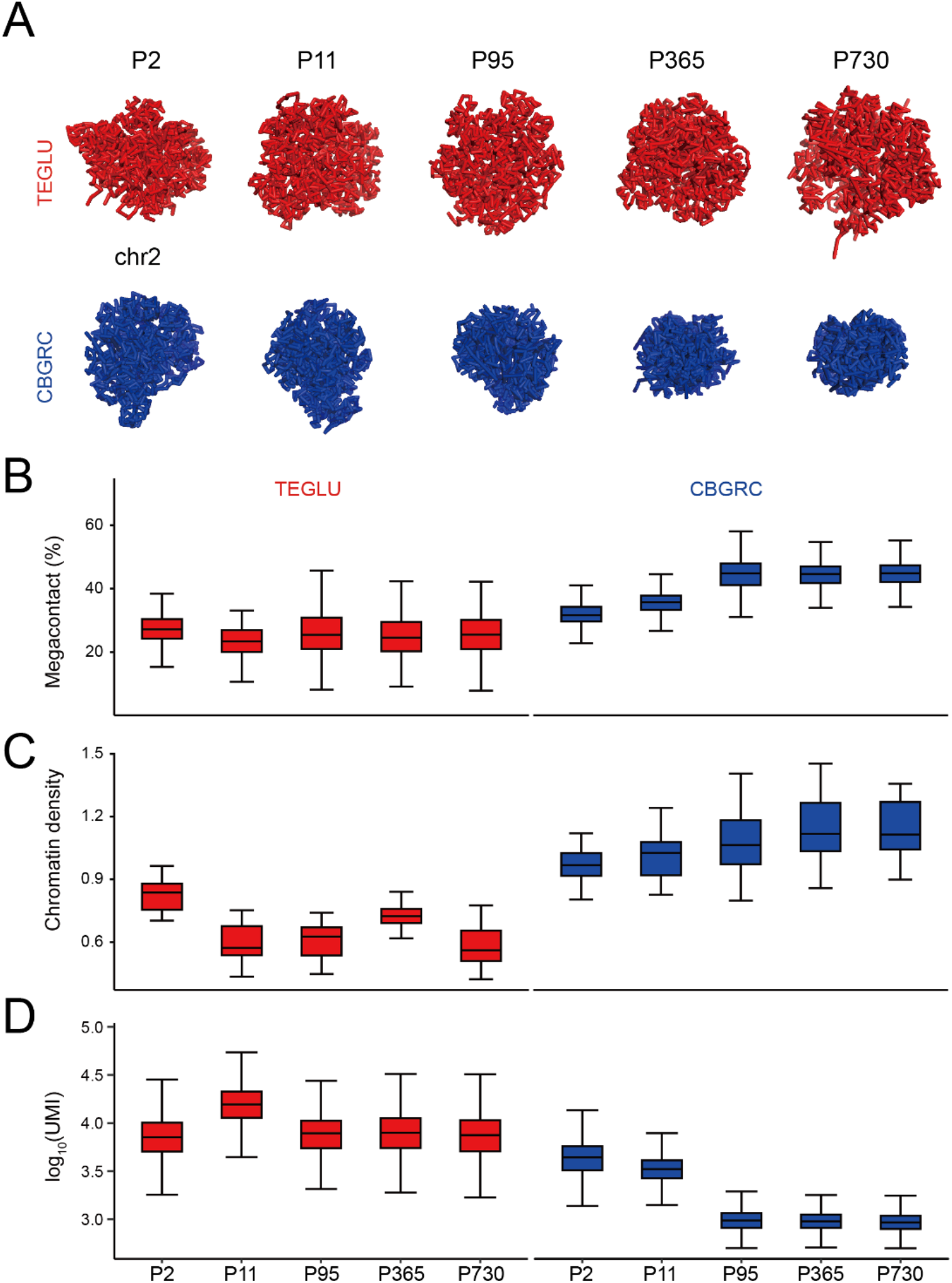
Nuclear volume dynamics of ExNs during aging. (**A**) Example view of 3D models of nuclear architectures from TEGLU and CBGRC across 5 age points. (**B**-**D**) Measurements of megacontact percentage (**B**), chromatin density inferred from the 3D models of nuclear architectures (**C**), and the global transcription activity (**D**) in ChAIR data.

## Notes

### Competing Interest Statement

The authors have declared no competing interest.

